# Defective cytoskeletal dynamics underlies the essential role of MRTF-SRF in IL-2 delivery to CD8^+^ T cells during infectious challenge

**DOI:** 10.1101/2023.08.26.554941

**Authors:** Diane Maurice, Patrick Costello, Francesco Gualdrini, Bruno Frederico, Richard Treisman

**Affiliations:** Signalling and transcription Laboratory, Francis Crick Institute, 1 Midland Road, London NW1 1AT, UK; Immunobiology Laboratory, Francis Crick Institute, 1 Midland Road, London NW1 1AT, UK; European Institute of Oncology (IEO), Istituto di Ricovero e Cura a Carattere Scientifico (IRCCS), Milan 20139, Italy

**Keywords:** CD8, LM-OVA, SRF, MRTF, IL-2, cytokines, actin, homotypic clustering, paracrine signaling

## Abstract

Paracrine IL-2 signalling underpins late primary CD8+ T cell expansion and differentiation that allow protection against viral infections, yet the requirements for effective delivery of IL-2 to recipient cells remain poorly understood. We show that the SRF transcription factor, a master regulator of cytoskeletal dynamics, is essential for the response to *L. monocytogenes* infection. SRF acts cell-autonomously with its actin-regulated MRTF cofactors *Mrtfa* and *Mrtfb* to sustain CD8^+^ effector T cell expansion and persistence of memory cells. MRTF-SRF activity is not required for initial TCR-mediated CD8^+^ T cell proliferation, but is necessary for subsequent IL-2 dependent expansion. Following TCR activation *in vitro*, *Mrtfab*-null CD8^+^ T cells produce IL-2 normally, but exhibit defective paracrine IL-2 signalling. Cluster formation by activated *Mrtfab*-null CD8^+^ T cells is impaired: clusters are smaller and less dense, have substantially reduced F-actin content, retain less IL-2, and exhibit defective cytoskeletal gene expression. Activated *Mrtfab*-null CD8^+^ T cells also exhibit defective homotypic clustering *in vivo*. The requirement for MRTF-SRF signalling for CD8^+^ T cell proliferation during infection thus reflects its involvement in cytoskeletal dynamics.

## INTRODUCTION

During infection, regulation of CD8^+^ T cell migration, proliferation and differentiation plays critical roles in successful pathogen clearance and establishment of protective immunity^1–3^. Upon interaction of naïve T cells with antigen-presenting cells (APCs), formation of the immune synapse facilitates TCR activation by peptide-MHC complexes and co-stimulatory adhesive signalling^4, 5^. Activated CD8^+^ T cells differentiate into highly proliferative short-lived effector cells (SLECs), which mediate pathogen clearance and undergo subsequent apoptosis, and a smaller population of memory precursor cells (MPEC) which give rise to self-renewing memory cells^6–8^. Strong TCR signals favour SLEC differentiation^8, 9^, while inactivation of mTOR favours MPEC formation and memory cell generation^10^.

While the initial proliferative response of CD8^+^ T cells is TCR-dependent, their subsequent proliferation and differentiation is modulated by IL-2 and other cytokines^11–14^ (see ^15, 16^ for review). The strength of IL-2 signalling controls the balance between effector and memory cell formation^17–, 19^, and the cytokine signal transducer STAT5 is essential for effector cell expansion^20, 21^. Both IL-2 and the transcription factor IRF4 are dispensable for early CD8^+^ T cell proliferation but necessary for sustained expansion and SLEC proliferation^11, 18, 22–24^. Effective cytokine delivery and activity is dependent on hetero- and homotypic T cell interactions^14, 25–27^. In keeping with this, SLEC expansion is dependent on the T cell integrin LFA-1 (αLβ2; CD11a/CD18)^27–29^, its ligand ICAM^27^, and its effector kinase PYK2^28^.

The SRF transcription factor controls receptor-regulated and cytoskeletal gene expression^30–32^ through interaction with two families of signal-responsive regulatory cofactors^33, 34^. The ERK-regulated Ternary Complex Factors (TCFs) control classical immediate-early genes associated with cell proliferation, while the Myocardin-Related Transcription Factors (MRTFs), which sense cellular G-actin concentration, control dozens of genes required for effective cytoskeletal dynamics^35–38^. During development, TCF-SRF signalling is essential for thymocyte positive selection^39–42^, while MRTF-SRF signalling is essential for seeding of HSC/P in the bone marrow^43^ and for thymocyte migration (PC, DM and RT, in preparation).

Here we demonstrate that MRTF-SRF signalling is also required for effective CD8^+^ T cell proliferation in response to *L. monocytogenes* infection. MRTF-SRF signalling is dispensable for the initial proliferative response to TCR activation, but required for the subsequent sustained expansion of activated CD8^+^ T cells in response to endogenous IL-2. *Mrtfab*-null T cells exhibited defective homotypic T-cell clustering, polarisation and motility *in vitro* and *in vivo*, and reduced basal and IL-2 induced cytoskeletal target transcription. Our findings demonstrate the importance of MRTF-SRF-dependent cytoskeletal dynamics and homotypic CD8^+^ T cell interactions for IL-2 induced proliferation.

## RESULTS

### SRF is essential for peripheral T cell expansion upon Listeria infection

To study CD8^+^ T-cell differentiation during infection, we used the well-characterised *L. monocytogenes*-OVA (LM-OVA) infection model^44, 45^. SRF is essential for T cell development, so to study its role in peripheral T-cell function we exploited a tamoxifen-inducible conditional *Srf* allele coupled with bone marrow reconstitution (Fig.1A). Bone marrow from *Srf*^f/f^ or *Srf*^+/+^ (WT) animals ubiquitously expressing a tamoxifen-regulated Cre derivative and the OVA-specific OT-I TCR (OT-I *Srf*^f/f^ TamCre; CD45.2, and OT-I WT TamCre; CD45.1) was used to reconstitute RAG-2^-/-^ mice. Six weeks later, mice were fed with tamoxifen for 15 days to inactivate *Srf* and allow SRF protein depletion in hematopoietic tissues (Fig.1B,S1A). The resulting OT-I WT TamCre and OT-I *Srf*^-/-^ TamCre CD8 T-cells were predominantly naïve, as assessed by CD62L, CD44, CD5, and CCR7 expression (Fig.S1B), and their failure to produce cytokines upon OVA peptide stimulation *in vitro* (Fig.S1C).

**Figure 1.**
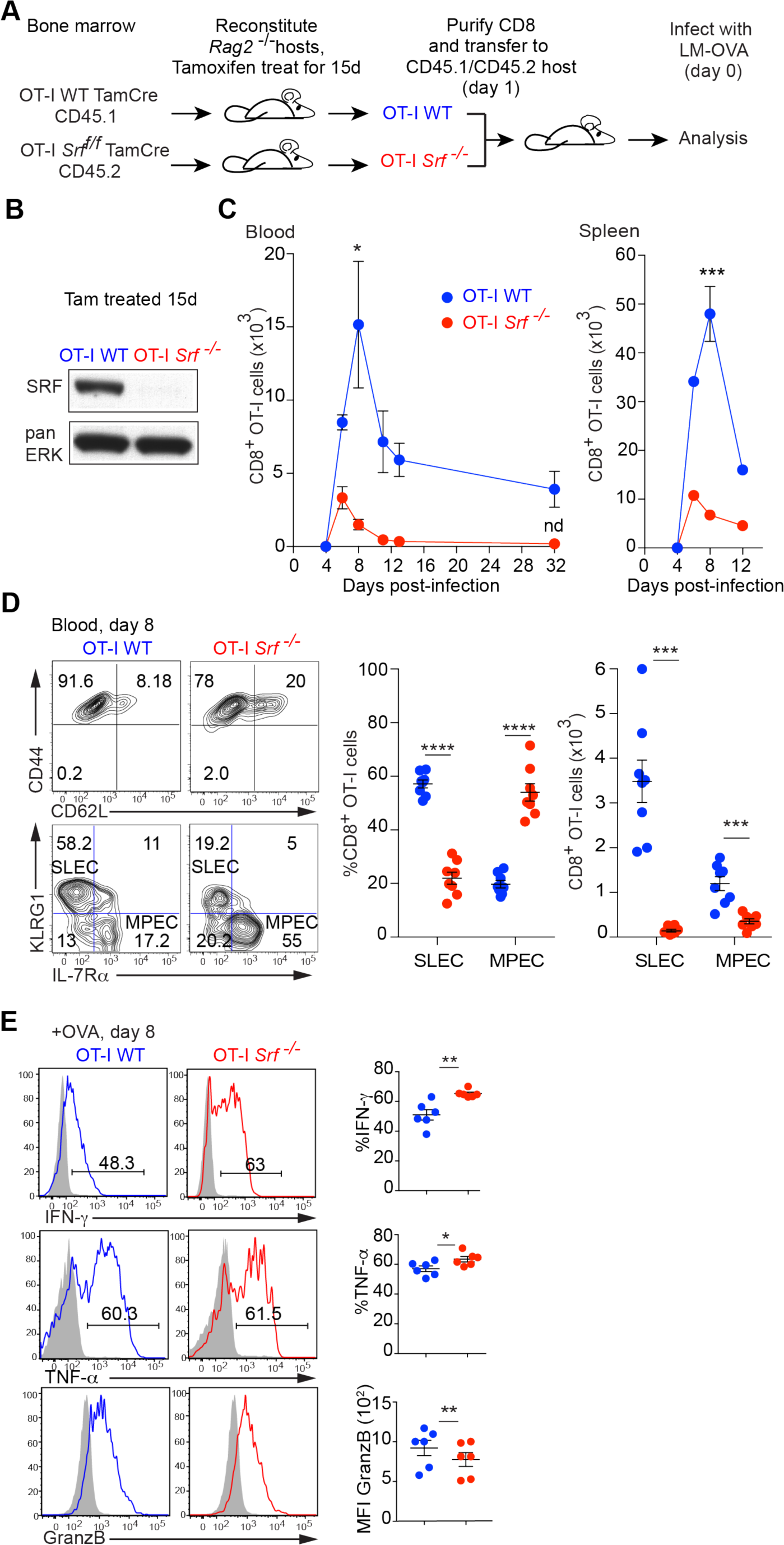
*Srf* is essential for the CD8^+^ T cell response to *Listeria* infection. **(A)** Experimental protocol. Irradiated *Rag2^-/-^* mice were reconstituted with bone marrow from OT-I WT TamCre or OT-I *Srf*^f/f^ TamCre mice and the resulting chimeras fed with tamoxifen for 15 days. Purified OT-I WT (CD45.1) and OT-I *Srf*^-/-^(CD45.2) naïve CD8^+^ T-cells were then co-transferred 1:1 into CD45.1/CD45.2 recipients, and infected with rLM-OVA the following day. **(B)** Immunoblot analysis of SRF protein in *Rag2^-/-^* reconstituted mice following 15 days tamoxifen treatment reveals complete *Srf* inactivation in OT-I *Srf*^-/-^cells. **(C)** Flow cytometry analysis following adoptive co-transfer of 5000 each OT-I WT and OT-I *Srf^-/-^* CD8^+^T cells upon rLM-OVA infection. Data are mean numbers ± SEM in blood (20μl) or spleen, 3 mice per time point, and are representative of ≥3 independent experiments; nd, not detectable. **(D)** Cell surface expression of activation markers CD44 and CD62L, and MPEC/SLEC markers KLRG1 and IL-7Rα on OT-I WT TamCre (CD45.1 gated) and OT-I *Srf*^-/-^ TamCre (CD45.2 gated) at day 8 post-infection in blood. Right panel, Proportions and numbers of SLEC (KLRG1^hi^, IL-7Rα^low^) and MPEC (KLRG1^low^, IL-7Rα^hi^) are shown. Data are means ± SEM; data points are individual mice, and are representative of ≥3 independent experiments. **(E)** CD8^+^ T cells were isolated 8 days post-infection, activated with 10nM SIINFEKL OVA peptide for 5h, and stained for IFN-γ, TNF-α and GranzB. Grey isotype control; black, experimental sample. Data are mean values ± SEM; data points represent individual mice.

To compare the responses of wildtype and *Srf*-null CD8^+^ T-cells to LM-OVA infection in the same environment we used an adoptive cotransfer strategy. OT-I WT and OT-I *Srf^-/-^* CD8^+^ T-cells were combined at 1:1 ratio, and 10,000 cells co-transferred into congenic C57BL/6 CD45.1/2 recipients, which were challenged 24h later with a priming dose of LM-OVA. Expansion of OT-I *Srf*^-/-^ effector CD8^+^ T-cells was greatly reduced in blood and spleen, and maximal at day 6 (Fig.1C). At day 8, the peak of wildtype CD8^+^ T cell expansion, both OT-I WT and OT-I *Srf^-/-^* CD8^+^ T-cells expressed CD44 similarly, indicating that activation was intact. Upon *ex vivo* stimulation with OT-I specific OVA peptide SIINFEKL, OT-I *Srf^-/-^*T-cells produced increased amounts of TNF-α and IFN-γ compared with OT-I WT cells, but granzyme B expression was slighlty decreased, suggesting their differentiation was affected (Fig.1E). Both CD62L and IL-7Rα were initially downregulated normally in OT-I *Srf^-^*^/-^ CD8^+^ T-cells, but by day 8 an enhanced proportion had re-acquired CD62L and IL-7Rα expression (Fig.S1D). Consistent with this, the OT-I *Srf^-/-^* CD8^+^ population displayed a decreased SLEC:MPEC ratio^6, 7^, reflecting a dramatic reduction in SLEC numbers (Fig.1D). By day 32 post-infection, when fully differentiated memory CD8^+^ T cells should be present, however, OT-I *Srf*^-/-^ CD8^+^ T-cell numbers had declined below the limit of detection (Fig.1C). During LM-OVA infection, SRF thus plays an essential and cell-autonomous role in robust CD8^+^ T-cell expansion and in generation and persistence of memory cells.

### SRF is required for CD8^+^ T memory cell expansion

To generate memory CD8^+^ T cells lacking SRF, we used the R26TamCre system to inactivate *Srf* once the response to initial infection was under way. OT-I WT and OT-I Srf^f/f^ cells were adoptively transferred to wildtype mice at 1:1 ratio, and the recipients treated with tamoxifen 4 days after LM-OVA infection (Fig.2A). In this setting, SRF was effectively depleted by day 6 post-infection (Fig.2B). Expansion of these OT-I “*Srf*^-/-^(post)” cells in blood was maximal at day 8: their SLEC-MPEC ratio was substantially unaltered (Fig.S2A), and they produced IFN-γ at comparable levels to OT-I WT cells upon activation with OVA peptide (Fig.S2B).

**Figure 2.**
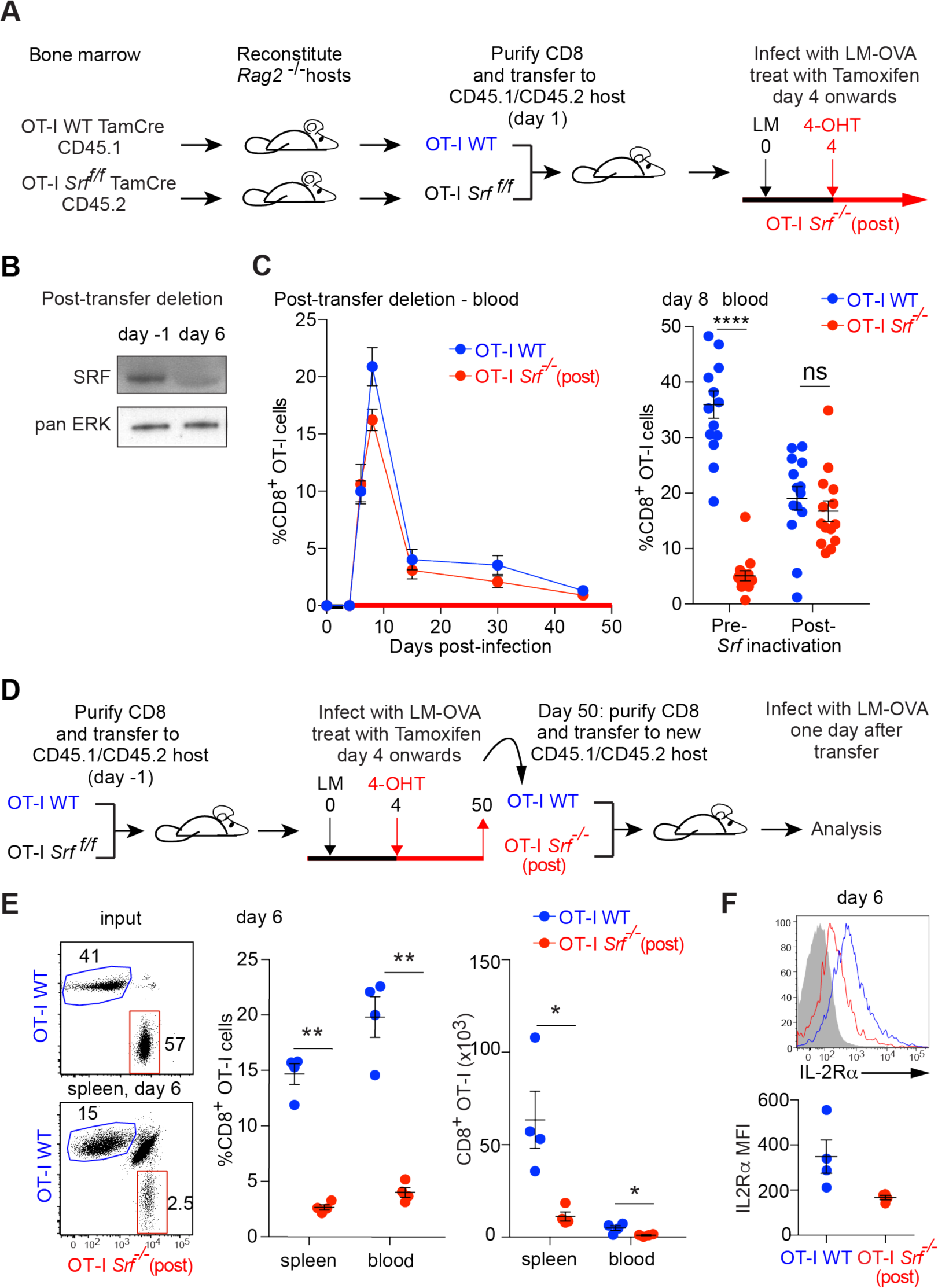
*Srf*-null memory cells cannot mount an effective recall response. **(A)** Protocol for post-infection inactivation of *Srf*. For post-infection, WT OT-I TamCre (CD45.1) and OT-I *Srf^f/f^* TamCre (CD45.2) CD8^+^ T cells were co-injected at a ratio 1:1 into CD45.1/CD45.2 hosts, which were infected with rLM-OVA the next day. OT-I *Srf*^-/-^(post): four days after infection, animals were fed tamoxifen to induce *Srf* inactivation. **(B)** Immunoblotting analysis of *Srf* following tamoxifen administration. **(C)** Expansion of WT OT-I TamCre and OT-I *Srf*^-/-^(post) CD8^+^ T cells in blood. Data show mean values ± SEM, 3 mice per time point. Representative of two independent experiments. Right, percentages of OT-I WT and OT-I *Srf*^-/-^ CD8^+^ cells at day 8 after infection in the blood of animals. OT-I *Srf*^-/-^(post): four days after infection, animals were fed tamoxifen to induce *Srf* inactivation. OT-I *Srf*^-/-^(pre): animals transferred with *Srf*-null cells as in Fig.1A. Data from 3 independent experiments, one animal per data point. **(D)** A schematic of OT-I WT (CD45.1) and OT-I *Srf*^-/-^(post) (CD45.2) memory cell populations purified at day 50 after primary LM-OVA infection from spleen of animals treated with the post-infection deletion regime as in Fig.2A and transferred at 1:1 ratio to a secondary CD45.1/CD45.2 host, which was subjected to infection the following day. **(E)** Expansion profiles, percentages and absolute cell counts of OT-I WT and OT-I *Srf^-/-^*(post) memory cells are compared 6 days after infection. Data show mean values ± SEM; data points represent individual mice. Statistical significance: paired t test. **(F)** IL-2Rα cell surface expression in OT-I WT (blue) and OT-I *Srf*^-/-^(post) (red) cells 6 days after memory immune response, performed as in (D).

OT-I *Srf*^-/-^(post) cells contracted with similar kinetics to OT-I WT cells, and putative *Srf*-null memory cells were readily detectable at day 45 (Fig.2C;Fig. S2C, S2D). These expressed elevated levels of the Tcm markers CD62L, CCR7 and IL-2Rβ, suggesting that late inactivation of *Srf* might enhance acquisition of central memory characteristics (Fig.S2C). Memory OT-I WT and *Srf*^-/-^(post) cells, defined by CD62L, CCR7 and IL-2Rβ expression, were purified and co-transferred at 1:1 ratio into new recipients (Fig.2D). The transferred *Srf*^-/-^(post) memory cells exhibited greatly reduced expansion upon LM-OVA challenge: at day 6 after infection, when the wildtype recall response reached its peak, the OT-I *Srf*^-/-^(post) numbers were only 20% that of wildtype (Fig.2E). Moreover, the re-transferred OT-I *Srf*^-/-^(post) CD8^+^ memory cell population did not maintain expression of IL-2Rα at day 6 following infection (Fig.2F). Thus, *Srf* is essential not only for the primary CD8^+^ T cell expansion but for reactivation and expansion of CD8^+^ T memory cells in response to infectious challenge.

### Requirement for SRF reflects the activity of its MRTF cofactors

We next used the co-transfer approach to investigate the contributions of the two families of signal-regulated SRF cofactors, the TCFs and the MRTFs^34^, to the CD8^+^ T-cell response during LM-OVA infection. SAP-1, encoded by *Elk4*, is the most abundant TCF in T cells^40, 41^, so we examined the response of animals co-transferred with OT-I WT and OT-I *Elk4*^-/-^ CD8^+^ T cells to LM-OVA infection (Fig.3A). Inactivation of *Elk4* affected neither the CD8^+^ T cell proliferative response (Fig.3B) nor the balance between SLECs and MPECs following infection (Fig.3C).

**Figure 3.**
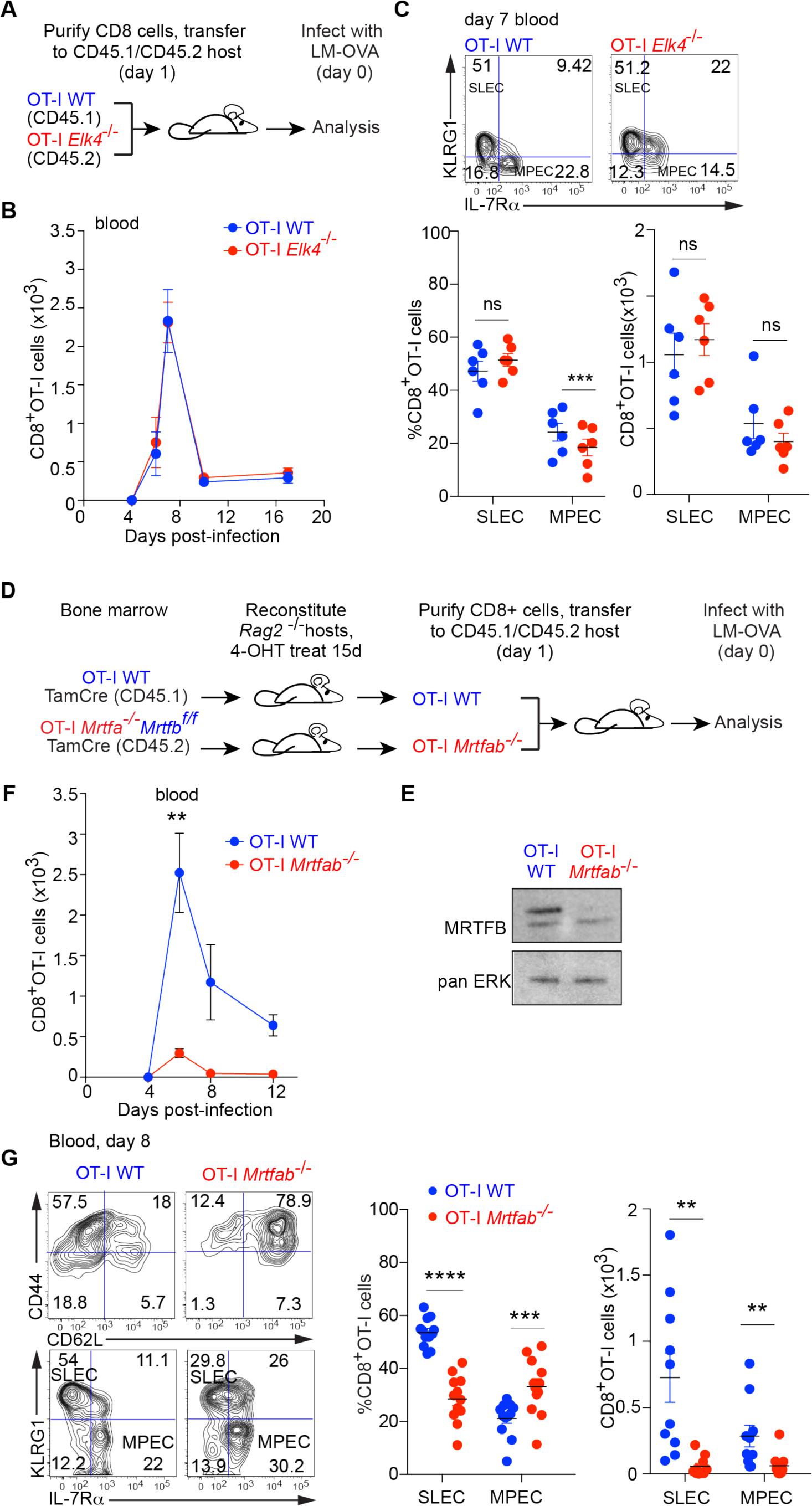
The MRTFs mediate signalling to SRF in the CD8^+^ T cell immune response. **(A)** Experimental protocol for analysis of OT-I *Elk4*^-/-^ CD8^+^ T cells. **(B)** Analysis by flow cytometry of expansion of 5000 adoptively co-transferred OT-I *Elk4*^-/-^ and OT-I WT CD8^+^ cells after infection, mean value ± SEM (n=6 mice). **(C)** Top, representative profile of cell surface expression of KLRG1 and IL-7Rα, 7 days post-infection; bottom, quantification of SLEC and MPEC. Data are mean values ± SEM; data points represent individual mice. **(D)** Irradiated *Rag2^-/-^* mice were reconstituted with bone marrow from OT-I WT TamCre or OT-I *Mrtfa^-^*^/-^*Mrtfb*^f/f^ TamCre mice. Following reconstitution, chimeras were treated for 15 days with tamoxifen and OT-I WT (CD45.1) and OT-I *Mrtfab*^-/-^ (CD45.2) naïve CD8^+^ T cells were purified and co-transferred 1:1 into CD45.1/CD45.2 recipients, followed by rLM-OVA infection the next day. **(E)** Immunoblot analysis before transfer reveals complete *Mrtfb* inactivation. **(F)** Analysis by flow cytometry of expansion of 5000 adoptively co-transferred OT-I WT and OT-I *Mrtfab*^-/-^ CD8^+^ T cells following infection. Data show mean numbers ± SEM in the blood (20μl). Data are representative of 2 independent experiments with 5 mice each. **(G)** Cell surface expression of CD44, CD62L, KLRG1 and IL-7Rα by flow cytometry as in Fig1D. Quantification of SLEC and MPEC. Data show mean values ± SEM; data points represent individual mice pooled from 2 independent experiments with 6 mice each.

To investigate the role of the MRTFs, we used a similar deletion strategy to that used for SRF, exploiting the viability of *Mrtfa*-null mice in conjunction with the conditional *Mrtfb*^fl/fl^ allele^46^. RAG-2 deficient hosts were reconstituted with OT-I WT TamCre CD45.1 or OT-I *Mrtfa*^-/-^*Mrtfb*^fl/fl^ TamCre CD45.2 bone marrow, treated with tamoxifen to inactivate *Mrtfb*, and subsequently co-transferred with OT-I WT cells to new hosts for infection analysis (Fig.3D,3E). The transferred OT-I *Mrtfab^-/-^* CD8^+^ T-cells were naïve, as assessed by CD62L and CD44 expression (Fig.S3A). As with Srf inactivation, by day 8 an enhanced proportion of Mrtfab-null cells had re-acquired CD62L and IL-7Rα expression (Fig.S3B). Upon LM-OVA infection, expansion of OT-I *Mrtfab*^-/-^ CD8^+^ T-cells was substantially attenuated (Fig.3F), and the SLEC/MPEC ratio was markedly reduced, with greatly impaired SLEC accumulation (Fig.3G). The requirement for SRF in CD8^+^ T-cell expansion following LM-OVA infection thus reflects the activity of its MRTF cofactors.

### Sustained proliferation requires MRTF-SRF signalling

To gain insight into the nature of the expansion defect in *Srf*-null T-cells, we examined the initial stages of infection in the spleen, the primary site of *L. monocytogenes* infection. To enable cell tracking during the early stages of infection, we used larger numbers of OT-I WT and OT-I *Srf^-/-^* cells for adoptive transfer (1-2 x 10^6^ each genotype). Following LM-OVA infection, both OT-I WT and OT-I-*Srf^-/-^* cell populations increased similarly until day 3 post infection, but OT-I *Srf*^-/-^ cells exhibited significantly reduced accumulation at day 4 (Fig.4A). A similar result was obtained with OT-I *Mrtfab^-/-^* cells (Fig.4B), which localised to the T-cell zones of the spleen, and upregulated the IL-2 receptor normally following LM-OVA infection (Fig.S4A,S4B). These expansion defects likely reflect decreased proliferation rather than increased cell death, since BrdU incorporation by OT-I *Srf*^-/-^ cells subsided after day 3 post-infection, while the amounts of active caspase-3 were similar to those in wildtype cells (Fig.4C). These observations suggest that initial TCR activation remains intact in OT-I *Srf^-^*^/-^ and OT-I *Mrtfab^-/-^* cells; indeed, naïve cells proliferated if anything more effectively than wildtype following activation *in vitro* by plate-bound anti-CD3/CD28 or OVA peptides (Fig.4D,4E; Fig.S4C; see Discussion). Taken together, these results show that MRTF-SRF signalling is required for effective CD8^+^ T cell proliferation at a step subsequent to initial T cell activation.

**Figure 4.**
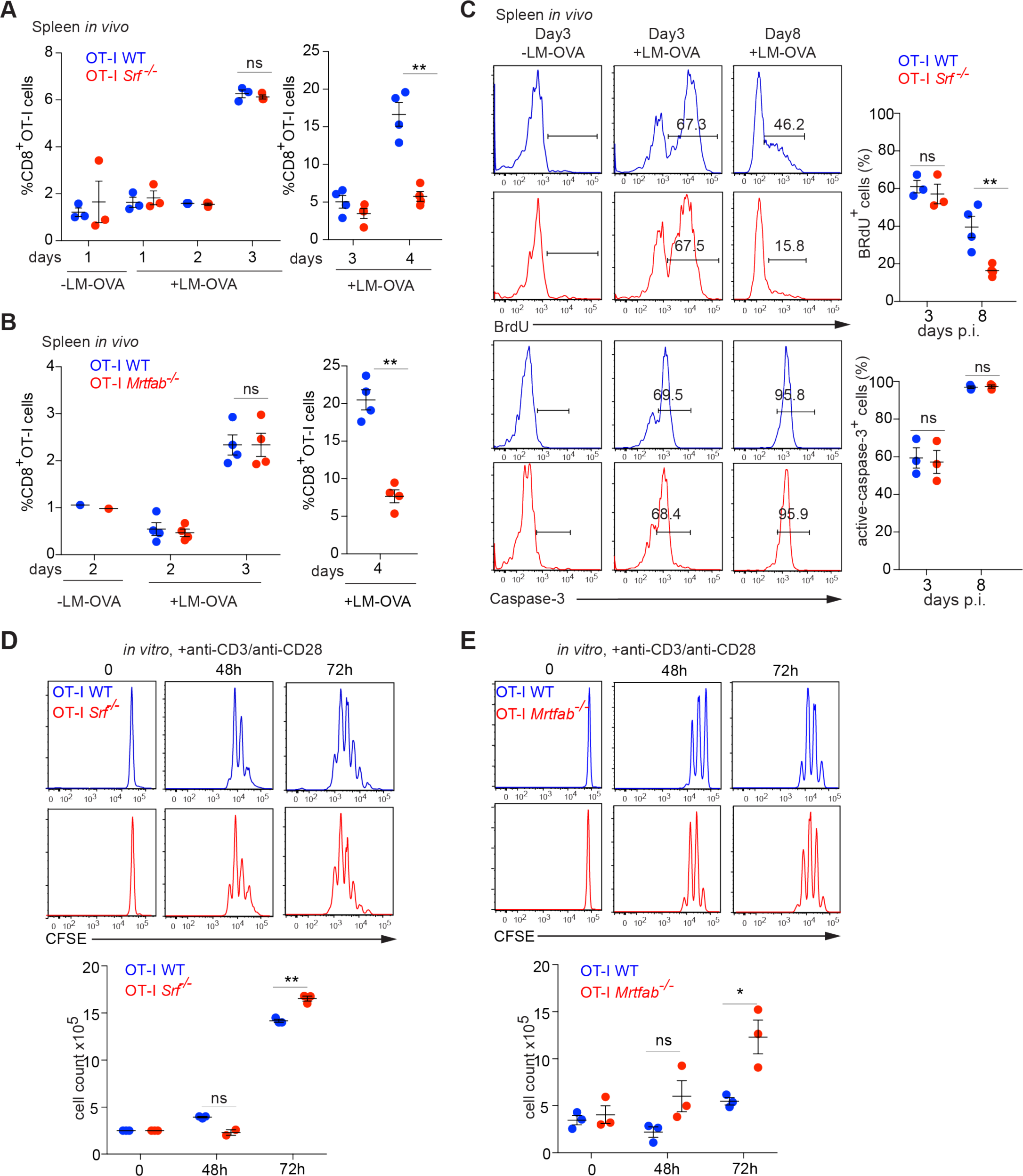
*Srf* is required for the sustained proliferation of CD8^+^ T cells. **(A)** Frequencies of adoptively co-transferred (1×10^6^ cells) OT-I WT TamCre and OT-I *Srf*^-/-^ TamCre in spleen from day 1 to day 4 p.i. Data show mean values ± SEM; data points represent individual mice. Representative of ≥2 experiments. **(B)** Frequencies of adoptively co-transferred (1×10^6^ cells) OT-I WT TamCre and OT-I *Mrtfab*^-/-^ TamCre in spleen from day 1 to day 4 p.i. Data show mean values ± SEM; data points represent individual mice. Representative of ≥2 experiments. **(C)** Upper panels, representative flow cytometry profiles of BrdU incorporation in OT-I WT and OT-I *Srf*^-/-^ in spleen labelled from days 1-3 and days 6-8 post-infection. Lower panels, representative flow cytometry profiles of caspase-3 activity in OT-I WT and OT-I *Srf*^-/-^ cells at day 3 and day 8 post-infection, Quantitation is as (A). **(D)** Representative CFSE division profiles of MACS-purified OT-I WT or OT-I *Srf^-^*^/-^ cells, either resting or activated with plate-bound anti-CD3/CD28 (5μg/ml) for the indicated times. Quantification of live cells, mean ± SEM. Data are representative of 3 independent experiments (each done in triplicate). **(E)** Representative CFSE division profiles of MACS-purified OT-I WT or OT-I*Mrtfab^-^*^/-^ cells, cultured as in (D).

### CD8^+^ T cell expansion failure is accompanied by IL-2 signalling deficits

The above results establish that following LM-OVA infection CD8^+^ T cells are activated and initiate proliferation, but expansion is not sustained in the absence of MRTF or SRF. We therefore investigated cell signalling during the transition to MRTF-SRF-dependent cell proliferation at days 3-4 post-infection. In both *Mrtfab*^-/-^ and *Srf*^-/-^ cells expression of IL-2Rα, the high affinity receptor for IL-2, which is induced both by TCR stimulation and by IL-2^13, 17, 19^ began to decline by day 3, before the onset of the expansion defect (Fig.5A,5B; Fig.S5A), and expression of IRF4, which is induced by TCR activation but also responsive to IL-2^22, 47^ behaved in a similar manner (Fig.5B; Fig.S5A).

**Figure 5.**
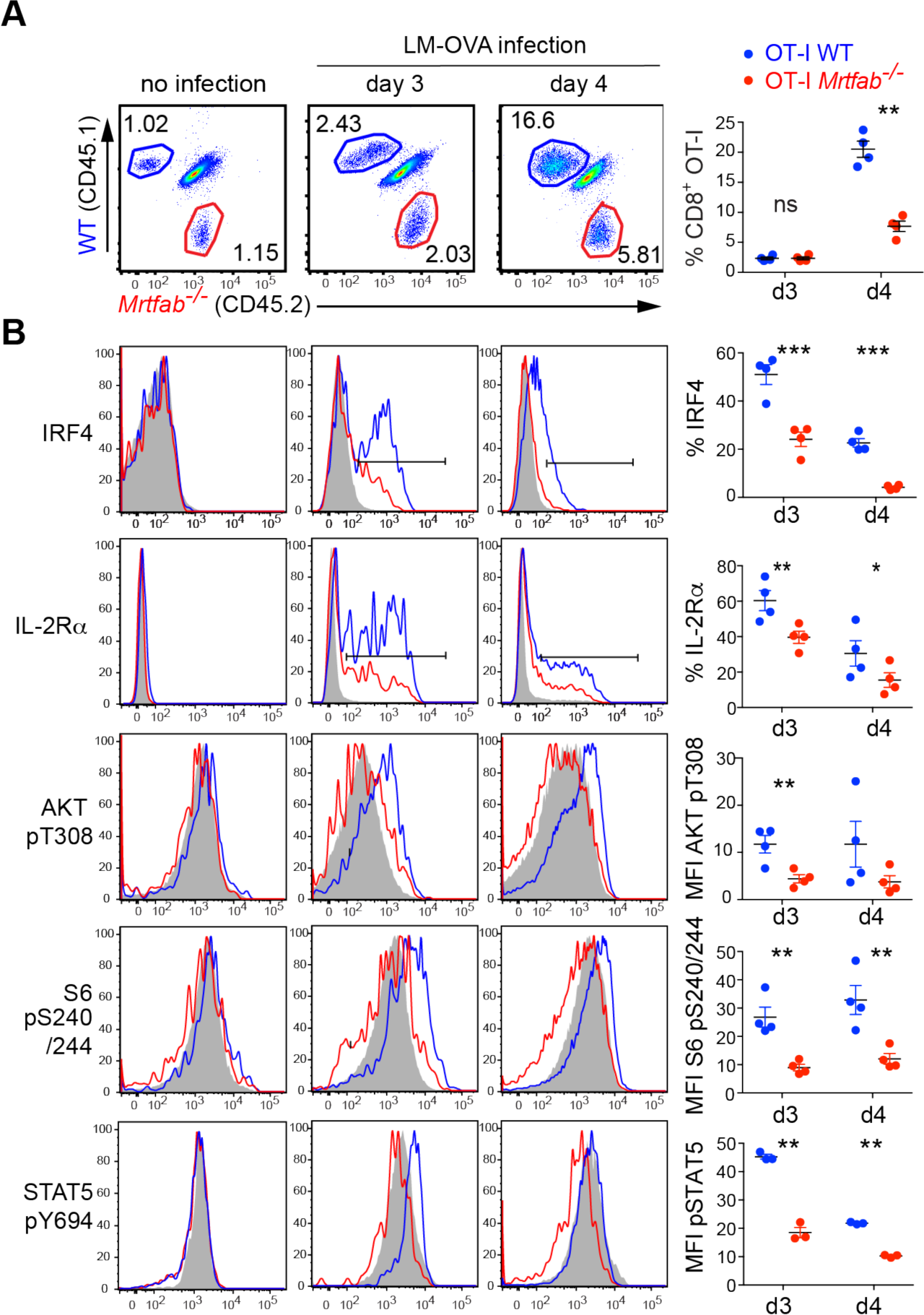
Expansion defects in *Mrtfab-* and *Srf-*null CD8^+^ cells are accompanied by defects in IL-2 signalling. **(A)** Flow cytometry analysis of splenocytes from mice co-transferred with OT-I WT CD45.1 and OT-I *Mrtfab*^-/-^ CD45.2 (1.10^6^ cells) and harvested at day3 or day4 p.i. Cells were stained with antibodies to CD8, CD45.1 and CD45.2 and CD8^+^ gated adoptive populations CD45.1 OT-I WT (blue) and CD45.2 OT-I *Mrtfab*^-/-^ (red) were analysed. **(B)** Relative levels of extracellular CD25 (IL-2Rα) protein expression, intracellular IRF4, pAKTthr308, pS6 240/244, pERK and pSTAT5 proteins from the populations in (A): OT-I WT (blue), OT-I *Mrtfab*^-/-^ (red) and endogenous CD45.1/CD45.2 CD8^+^ T cells (grey). Graphs show quantitations of protein expression by frequencies or MFI. Each data point represents a single mouse (n=4) in one representative experiment. Experiment has been done ≥3 times. Data show mean values ± SEM; Statistical significance was by paired t test.

In OT-I WT cells, activating phosphorylation of Akt T308 and phosphorylation of S6 S240/244 were readily detectable at day 3 (Fig.5B); phosphorylation of S6 was sensitive to the mTORC1 inhibitor rapamycin, consistent with the involvement of PI3K-Akt-mTOR signalling (Fig.S5B). In contrast, both OT-I *Srf*^-/-^ and OT-I *Mrtfab*^-/-^ CD8^+^ T cells exhibited reduced Akt T308 and S6 S240/244 and S235/236 phosphorylation, and the latter was not reduced by rapamycin treatment (Fig.S5A,S5B). Since IL-2 induces S6 phosphorylation via PI3K-Akt-mTOR signalling^48^, these results indicate that IL-2 signalling is defective in *Mrtfab*- and *Srf-*null CD8 T cells. Consistent with this view, STAT5a/b Y694 phosphorylation, a specific reporter of cytokine signalling (reviewed by ^16, 49^), was significantly reduced in OT-I *Mrtfab*^-/-^ CD8^+^ T cells (Fig.5B). These data suggest the failure to sustain CD8^+^ T cell expansion in *Mrtfab*- and *Srf-*null cells reflects a defect in IL-2 signalling.

### *Srf*^-/-^ and *Mrtfab*^-/-^ CD8^+^ T cells remain responsive to IL-2

We next tested whether the defect in IL-2 signalling lies at the level of signal receipt or response. *Mrtfab-* or *Srf*-null splenic T cells were harvested 3 days post-infection, when proliferation of OT-I WT and OT-I *Mrtfab*^-/-^ cells was comparable, and cultured for 72h either alone, with IL-2, or with IL-12, which potentiates IL-2Rα expression^50^. In the absence of added cytokine, OT-I WT cells expanded more than OT-I *Mrtfab*^-/-^ cells, and IL-2Rα expression declined to comparable levels on both populations (Fig.6A). Culture with IL-2 substantially increased both IL-2Rα expression and proliferation to comparable extents in wildtype and *Mrtfab*^-/-^ cells, while IL-12 restored IL-2Rα expression, but did not enhance proliferation (Fig.6A). Thus, MRTF-SRF signalling is not required for CD8^+^ cells to respond to IL-2 *in vitro*, or for IL-2Rα expression *per se*. Consistent with this, following activation and culture in IL-2, OT-I WT and OT-I *Srf^-/-^* cells purified from uninfected mice exhibited comparable abilities to kill OVA-pulsed EL4 target cells *in vitro*, as assessed by intracellular active caspase-3 staining (Fig.S6A).

**Figure 6.**
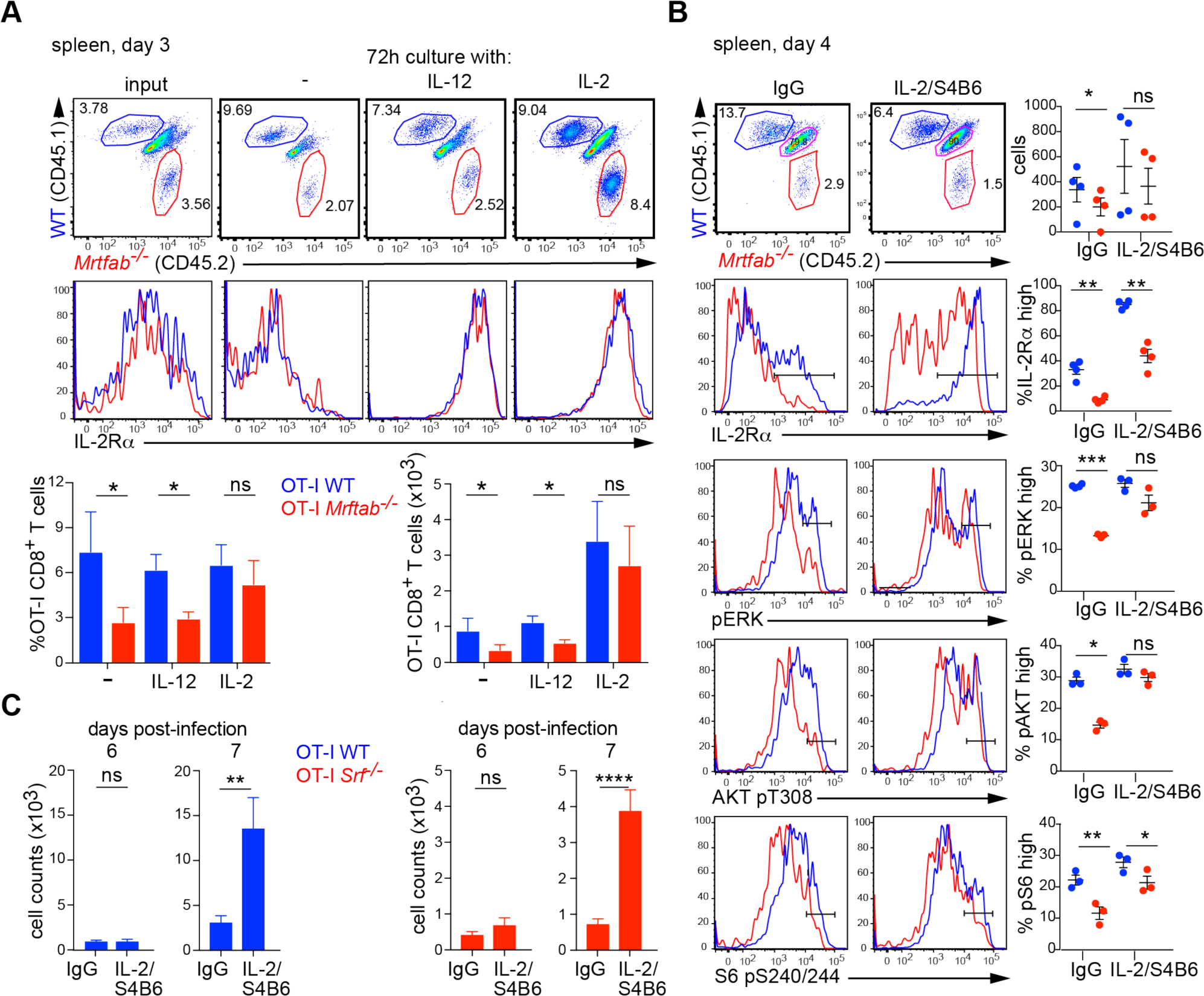
*Srf-* and *Mrtfab*-null cells remain responsive to exogenous IL-2. **(A)** Co-transferred OT-I WT and OT-I *Mrtfab*^-/-^ cells were harvested from spleen 3 days post LM-OVA infection (input) and cultured a further 3 days without cytokine, with IL-12 or IL-2. Top panels, dot-plots of CD8^+^ gated splenocytes, with histograms indicating the amount of IL-2Rα (CD25) expression. Bottom, relative proportions and numbers of OT-I WT and OT-I *Mrtfab*^-/-^ cells after 72h culture (mean ± SEM, n=3 mice). This is a representative experiment out of 2 independent experiments with ≥3 mice each. **(B)** Potentiation of OT-I *Mrtfab*^-/-^ signalling *in vivo* by administration of IL-2/S4B6 complexes. Mice were co-injected with 1x 10^6^ each OT-I WT and OT-I *Mrtfab*^-/-^ CD8^+^ T-cells and infected with Listeria the following day. Mice were treated with either IL-2/S4B6 complexes or control IgG from day 1 to 3 post-infection. Dot plots of CD8-gated splenocytes at day 4 post-infection show relative proportions of OT-I wildtype (CD45.1) and OT-I *Mrtfab*^-/-^ (CD45.2) cells. Histograms show IL-2Rα expression, pERK, AKT pT308 and S6 pS240/244. Quantitation is on the right (n=2, at least 3 mice per condition per experiment). **(C)** Potentiation of OT-I WT and OT-I *Srf*^-/-^ proliferation *in vivo* by administration of IL-2/S4B6 complexes from day 3 to 5 post-infection. Mice were injected with 5000 cells each OT-I WT and OT-I *Srf*^-/-^ CD8^+^ T-cells, infected with Listeria the following day, and cell numbers in the blood are quantified at 6 and 7 days after infection. Data are mean numbers ± SEM in blood (20μl), 10 mice per time point.

Next, we potentiated IL-2 signalling *in vivo* using IL-2/S4B6 antibody complexes, which prolong the half-life of IL-2, and signal through IL-2Rβ (CD122)^51^. IL-2/S4B6 injection from day 1 to day 3 post-infection potentiated IL-2Rα expression, ERK and AKT activation, and S6 phosphorylation in both OT-I WT and OT-I *Mrtfab*^-/-^ cells at day 4 and restored activation and proliferation of OT-I *Mrtfab*^-/-^ cells to levels comparable to wildtype (Fig.6B). Consistent with this, injection of IL-2/S4B6 complexes at days 3-5 post-infection substantially boosted the expansion of both OT-I WT and OT-I *Srf*^-/-^ cells (Fig.6C). Thus cells deficient in MRTF-SRF signalling remain sensitive to exogenously supplied IL-2.

### MRTF-dependent homotypic clustering is required for IL-2 signalling following TCR activation

The experiments in the preceding sections show that although *Mrtfab*-null and *Srf-*null CD8^+^ T cells retain the ability to respond to exogenous IL-2, they nevertheless exhibit defective ability to receive endogenous IL-2 signals, leading to a failure to maintain IL-2Rα expression and proliferative expansion after the priming phase. To investigate the basis of this defect, we first used an *in vitro* approach to assess how endogenous IL-2 signalling contributes to IL-2Rα expression following TCR activation. Wildtype, *Srf*-null and *Mrtfab*-null CD8^+^ T cells activated IL-2 gene transcription and produced IL-2 at comparable levels following activation by plate-bound anti-CD3/CD28 (Fig.S7A,S7B), and responded comparably to titration of exogenous IL-2 (Fig.S7C). At 24h following TCR activation, IL-2 blockade reduced IL-2Rα expression and abolished STAT5 Y694 phosphorylation, indicating that even at this early time IL-2 signalling is active (Fig.7A,7B). IL-2 blockade also inhibited the sustained expression of IRF4 at late times during continuous TCR activation (Fig.S7D). Although IL-2Rα expression remained essentially intact, OT-I *Mrtfab*^-/-^ cells exhibited decreased STAT5 Y694 phosphorylation, indicating a defect in IL-2 signal transduction (Fig.7A,7B).

**Figure 7.**
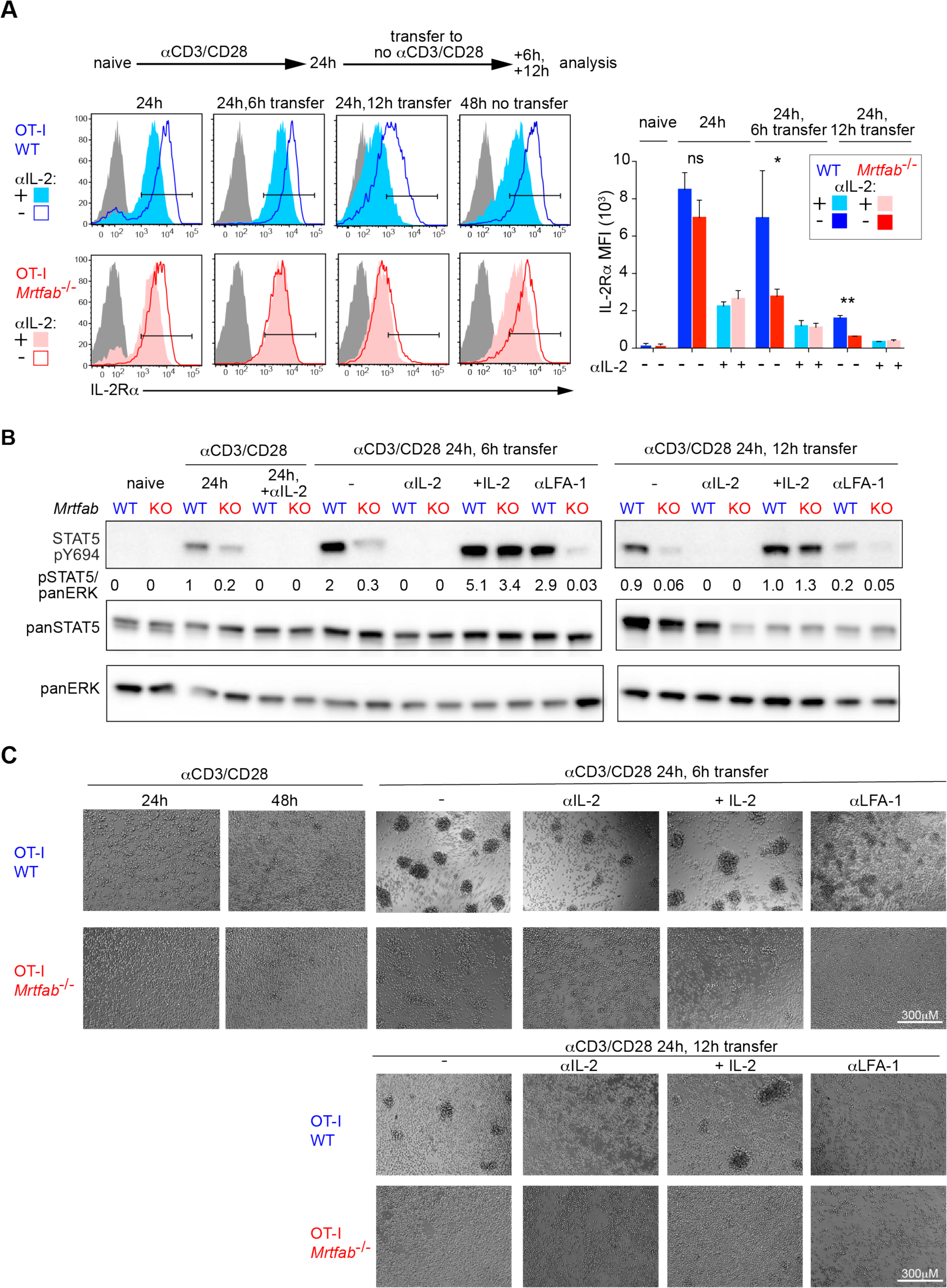
Deficient IL-2 signalling in activated *Mrtfab*^-/-^ OT-I cells correlates with defective homotypic clustering. **(A)** MACS purified OT-I WT and OT-I *Mrtfab*^-/-^ isolated from lymph nodes from tamoxifen-fed bone marrow-reconstituted mice are activated *in vitro* with plate-bound anti-CD3/anti-CD28 (5ug/ml) in the presence or absence of anti-IL2 blocking antiboby (JES6-1A12)(10μg/ml) for the times indicated. After 24h of TCR/CD28 activation, cells are transferred or not with supernatant to uncoated wells for an additional 6 or 12 hours of culture. Cells are harvested and stained for IL-2Rα and CD8 followed by fixation. Histograms of overlayed OT-I WT (blue) and OT-I *Mrtfab*^-/-^ (red) CD8^+^ cells are presented with quantification of IL-2Ra MFI +/-SEM for ≥3 independent experiments. **(B)** Stat5 phosphorylation (pSTAT5 Y694) measured in the presence or absence of anti-IL2 blocking antibody (JES6-1A12) (10μg/ml), ^r^mIL2(20ng/ml) or anti-CD11a/ anti-LFA-1 blocking antibody (10μg/ml) by western blot. Quantification below (pStat5/panErk). Data presented is representative of ≥3 biological replicates across all conditions. **(C)** Brightfield images of purified OT-I WT and OT-I *Mrtfab*^-/-^ generated and cultured as in (A) and (B). Images are representative of ≥3 independent experiments. Scale bar, 300μm.

These experiments show that both the TCR and endogenously-produced IL-2 contribute to IL-2Rα expression, even at early times of activation. To investigate signalling by endogenous IL-2 in the absence of continuing TCR activation, we activated cells for 24h and then transferred them to a new plate without anti-CD3/CD28. IL-2Rα expression in OT-I WT cells declined slowly following transfer, but declined much more rapidly in OT-I *Mrtfab*^-/-^ cells; upon IL-2 blockade the kinetics of IL-2Rα decline were similar in both genotypes, presumably reflecting declining TCR activity (Fig.7A). Signalling by the IL-2 receptor, as measured by STAT5 Y694 phosphorylation, initially increased upon transfer in OT-I WT, but not OT-I *Mrtfab^-/-^* cells, before decreasing in both at later times, and the increase was dependent on IL-2 (Fig.7B). Thus in this setting, continued expression of IL-2Rα is dependent both on the MRTFs and endogenous IL-2. The rapid down-regulation of IL-2Rα in *Mrtfab*-null cells upon cessation of TCR activation is therefore a consequence of defective IL-2 signalling rather than its cause.

Homotypic clustering plays an important role in T cell responses to cytokines^26, 27^, so we examined cell interactions in the transfer assay. Upon transfer to culture without anti-CD3/CD28, activated OT-I WT cells rapidly came together into large clusters, whose formation was substantially inhibited upon blockade of IL-2 or CD-11a (Fig.7C). In contrast, cluster formation by activated OT-I *Mrtfab*^-/-^ cells was greatly reduced, even though cell surface expression of LFA-1 and its ability to switch to high-affinity conformation upon TCR stimulation remained intact (Fig.S7E,F). Moreover, cluster formation by OT-I *Mrtfab*^-/-^ cells was not rescued by exogenous IL-2, in contrast to STAT5 phosphorylation (Fig.7C). Taken together these data are consistent with a model in which MRTF-SRF dependent clustering facilitates receipt of paracrine IL-2 signals.

### *Mrtfab*-null CD8^+^ T cells exhibit defective cytoskeletal and IL-2 induced gene expression

The results presented above show that while exogenous IL-2 can restore IL-2Rα expression and activity in *Mrtfab*-null CD8^+^ T cells, it does not restore clustering, suggesting that clustering is a prerequisite for effective endogenous IL-2 signalling. MRTF-SRF signalling plays a central role in cytoskeletal gene expression^33, 36^, so we next compared gene expression in wildtype and *Mrtfab*-null OT-I CD8^+^ T cells. Since activated wildtype cells, but not *Mrtfab*-null cells, can cluster and respond to endogenous IL-2, we compared gene expression profiles in activated cells with and without IL-2 stimulation. To evaluate the effects of TCR activation and IL-2 stimulation separately, naïve CD8^+^ T cells were activated by TCR crosslinking for 24h, then cultured for 16h without TCR crosslinking in the presence of IL-12 to maintain IL-2Rα expression (“TCR-activated/rested” cells), before stimulation with IL-2^52^ (Fig.8A).

**Figure 8.**
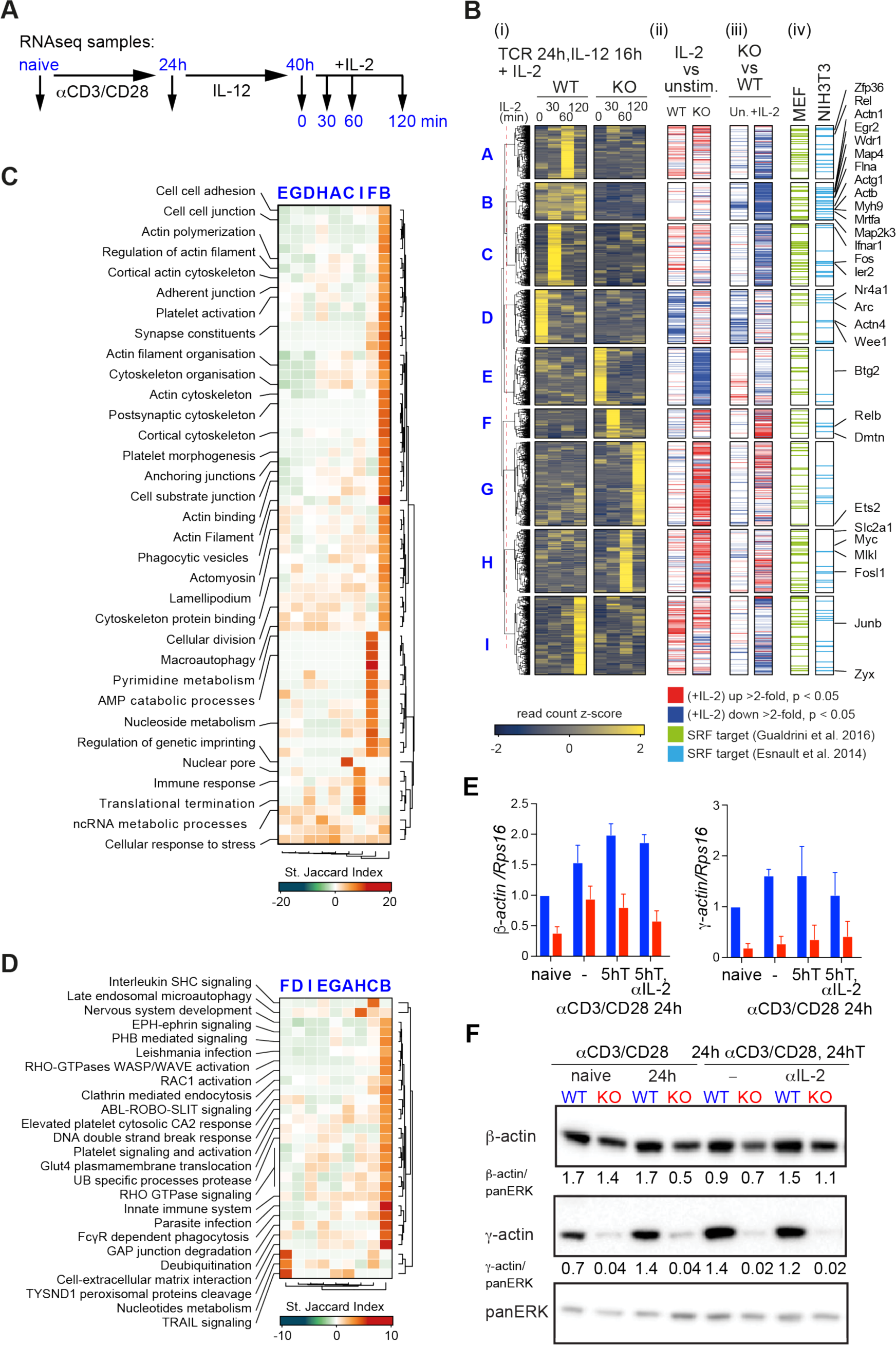
Gene expression deficits in OT-I *Mrtfab^-/-^* cells. **(A)**. Purified naïve wildtype or *Mrtfab*-null OT-I CD8^+^ T cells were stimulated as shown^52^. RNAseq was carried out on naïve cells, cells activated by TCR crosslinking for 24h, or activated cells cultured in IL-12 for 16h (“activated/rested cells”) and then stimulated with IL-2 for various times. Data show mean values ± SEM of three biological replicates. (B). (i) Normalized z-scored read counts of activated/rested cells stimulated with IL-2, grouped by unsupervised clustering. (ii) Genes showing significant changes upon IL-2 stimulation at any time point in wildtype (left) and *Mrtfab*-null cells (right). (iii) Genes whose expression is impaired by MRTF inactivation, in activated/rested cells (left) or IL-2 stimulated cells (right). (iv) Genes identified as candidate SRF targets in TPA-stimulated MEFs^37^ or serum-stimulated NIH3T3 fibroblasts^36^. **(C)** Gene ontology categories significantly overrepresented (P < 0.01) in the clusters identified in (B), summarised by standard Jaccard index. **(D)** Reactome pathway categories (https://reactome.org) significantly overrepresented (P < 0.01) in the clusters identified in (B). **(F)** Expression of β and γ actin in OT-I WT and OT-I *Mrtfab*^-/-^ cells following activation assessed by immunoblotting. MACS purified OT-I WT and OT-I *Mrtfab*^-/-^ cells isolated from spleen from tamoxifen-fed bone marrow reconstituted mice were activated *in vitro* with plate-bound anti-CD3/anti-CD28 (5ug/ml) for the indicated times. After 24h of TCR/CD28 activation, cells are transferred with supernatant to uncoated wells for an additional 24h of culture in the presence or absence of anti-IL2 blocking antibody (JES6-1A12). **(G)** Expression of β and γ actin in MACS purified OT-I WT and OT-I *Mrtfab*^-/-^ lymph nodes cells following TCR activation assessed by qRT-PCR. Cells are activated as in **(F)** but transferred and cultured for only 5h in the presence or absence of anti-IL-2 blocking antibody (JES6-1A12). Data show mean values ± SEM of three independent experiments with naïve OT-I-WT normalised to 1.

RNAseq analysis identified 3039 genes that exhibited significant differential expression in TCR-activated/rested *Mrtfab*-null cells compared to wildtype cells, regardless of IL-2 stimulation (Fig.8B, Table S1). These genes were grouped into 9 clusters (A-I) on the basis of their normalized z-scored read counts (see Materials and Methods). Four clusters (B,A,C, and I) encompassed 1134 genes whose transcript level was significantly reduced in *Mrtfab*-null cells (Fig.8B). Clusters A,C, and I represent IL-2 inducible genes, grouped according to induction kinetics. These were significantly enriched in gene ontology and reactome categories related to immune cell and rho GTPase signalling (Fig.8C,8D; Table S2; see Discussion). In contrast, the 200 genes in cluster B exhibited significantly impaired expression in TCR-activated/rested *Mrtfab*-null cells but were not significantly induced by IL-2. Cluster B is enriched in GO categories related to actin cytoskeletal regulation, and reactome signalling pathway components involved in rho-family GTPase signalling, infection, innate immunity, and immune cell biology (Fig.8C,8D; Table S2), and includes many MRTF-SRF targets^36, 37^. A further 1369 genes expressed at elevated levels in *Mrtfab*-null cells were significantly enriched in gene ontology categories involved in various metabolic processes (Fig.8B,8C, clusters E-H; Table S2).

To gain more insight into cluster B, we examined the relationship between gene expression in naïve cells and TCR-activated/rested cells prior to stimulation with IL-2. First, we used unsupervised clustering to identify groups of genes differentially responsive either to the initial 24h TCR activation (Fig.S8A,S8B, groups 1-11), or to subsequent resting in IL-12 for 16h (Fig.S8C, groups 1’-10’). We then used the Standardized Jaccard Index and hypergeometric testing to assess how these gene groups were related to genes that exhibited differential response to IL-2 stimulation in TCR-activated/rested cells. Cluster B genes exhibited a strong overlap with those genes whose strong response to initial TCR activation was abolished in *Mrtfab*-null *c*ells (cluster 10, Fig.S8B; Fig.S8D), and with those genes induced upon TCR activation and whose expression was maintained or increased upon resting in IL-12 (clusters 9’ and 10’, Fig.S8C, Fig.S8E).

Taken together, these results support a model in which the cytoskeletal defects associated with MRTF inactivation reflects defective expression of cytoskeletal genes and regulators accompanied by the de-regulation of IL-2 controlled genes. Indeed both naïve and activated OT-I *Mrtfab*^-/-^ cells exhibited significantly reduced levels of β and γ actin mRNA and protein (Fig.8E,8F; see Discussion).

### Abnormal *Mrtfab*-null CD8^+^ T cell clustering impairs IL-2 delivery

To study the role of the MRTFs in cluster formation in the absence of any confounding effects arising from TCR engagement or IL-2 signalling, we analysed clustering following T cell activation by PDBu and ionomycin. In this setting both OT-I WT and OT-I *Mrtfab^-/-^* cells upregulated IL-2Rα, LFA-1 and ICAM-1 similarly, but cluster formation by OT-I *Mrtfab*^-/-^ cells was impaired (Fig.S9A,S9B). OT-I *Mrtfab*^-/-^ clusters contained fewer, less polarised cells, were less densely packed, and contained almost 8-fold less F-actin (Fig.9A, 9B). In both genotypes PDBu/ionomycin-induced cluster formation was only weakly inhibited by IL-2 blockade but was dependent on LFA-1. (Fig.S9A,S9B). In spite of normal IL-2Rα expression in PDBu/ionomycin-induced *Mrtfab*-null CD8^+^ T cells, STAT5 pY694 phosphorylation was diminished, indicating that signalling by endogenously produced IL-2 is defective in this setting (Fig.S9C).

**Figure 9.**
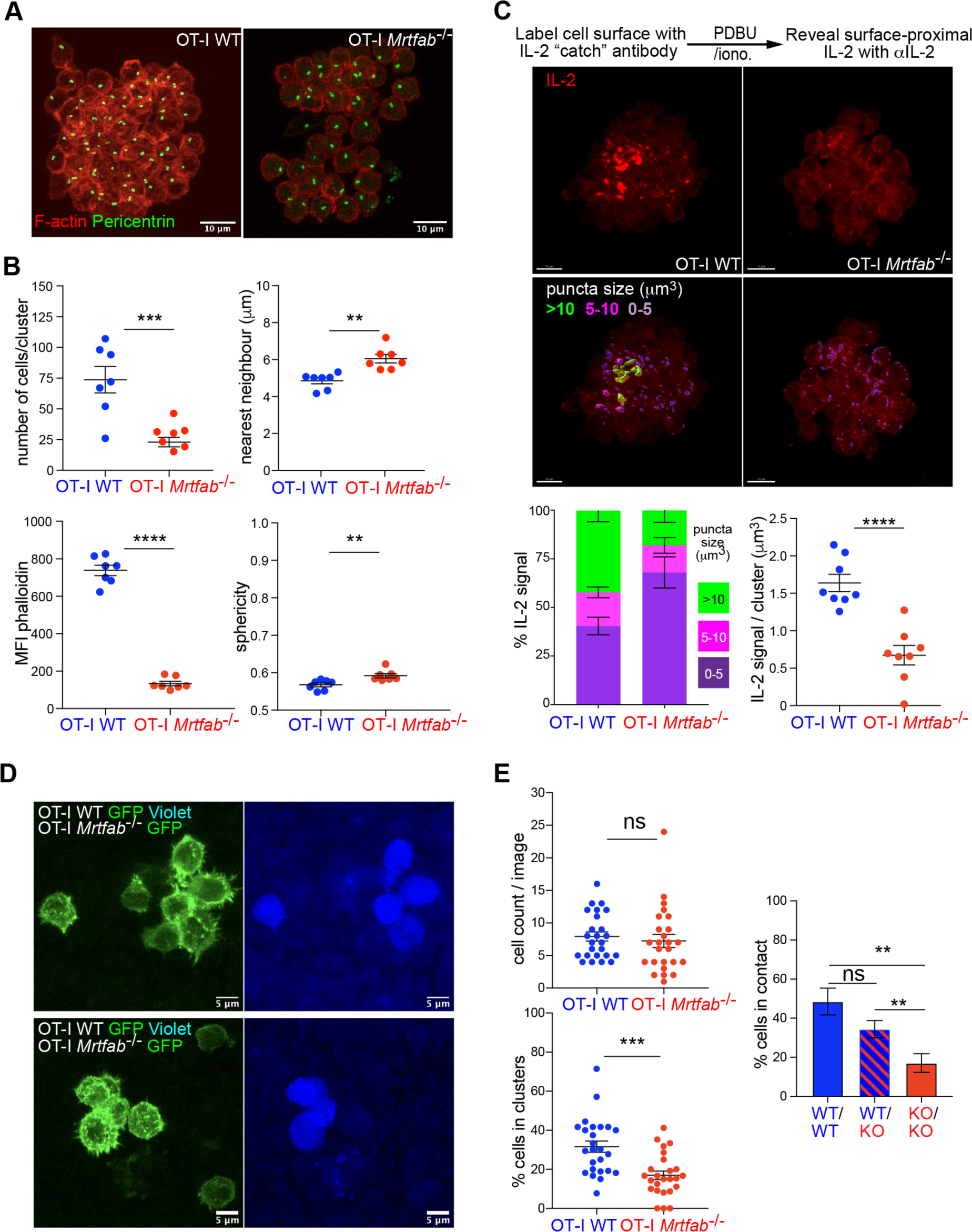
Defective clustering by OT-I *Mrtfab*^-/-^ cells *in vitro* and *in vivo*. **(A)** Defective clustering following 18h activation by PDBu/ionomycin *in vitro*. Representative confocal microscopy images of OT-I WT or OT-I *Mrtfab*^-/-^ clusters. Top, maximum intensity projections of Z stacks, bottom, single confocal XY plane. MACS-purified cells were stimulated for 18h with PDBu and Ionomycin to induce cluster formation. After fixation on poly-lysine coated slides, permeabilized clusters were stained with phalloidin, anti-pericentrin antibody and DAPI as indicated. **(B)** Characterisation of the clusters by cell number, distance to the nearest neighbour, F-actin content, and sphericity. Each data point is a determination for an individual cluster, shown as mean values ± SEM. Statistical significance, unpaired t test. **(C)** Visualisation of endogenous IL-2. Top, schematic of the catch assay. OT-I WT or OT-I *Mrtfab*^-/-^ cells were coated with IL-2 catch reagent, then stimulated with PDBu/ionomycin for 18h, and IL-2 retained within the cluster revealed using phycoerythrin-conjugated IL-2 antibody. Centre, representative images of cell clusters either stained for IL-2 (upper panels) or false-coloured according to IL-2 puncta size (lower panels). Bottom, size distribution of IL-2 puncta (left) and IL-2 signal volumes (right), determined for 8 clusters of each genotype in two independent experiments. Scale bar, 10μm. **(D)** *Mrtfab^-/-^* cells exhibit defective clustering *in vivo*. OT-I WT LifeactGFP (CellTrace violet-labelled) and OT-I *Mrtfab^-/-^* LifeactGFP were coinjected at 1:1 ratio i.v. into WT mice, immunised the next day with DNGR-1-OVA+anti-CD40 by subcutaneous injection into both hocks. The draining inguinal lymph nodes were isolated 24h later for confocal microscopy. Representative images with OT-I WT and OT-I *Mrtfab*^-/-^ LifeactGFP (green) (left) and OT-I-WT cell Trace violet (right). Regions containing clusters demonstrate the majority of the cells participating in clusters(≥ 2 cells) to be OT-I WTLifeactGFP. Note that expression of LifeactGFP is lower in *Mrtfab*-null cells because its expression is controlled by CAG promoter^70^, which is MRTF-dependent. **(E)** Quantitation of the experiment presented in (D). Top left, numbers of GFP+ cells present in each region analysed (top left) (n=25 images); bottom left, percentage of OT-I-WT or OT-I *Mrtfab*^-/-^cells in clusters out of total cells per image (n=25 images); dots represent data from each image. Right, mean frequencies of WT-WT, WT-KO and KO-KO contacts in two independent experiments. Data are mean values ± SEM; Statistical significance, paired t test.

We assessed whether MRTF activity affected presentation of IL-2 to cells within the cluster, using the IL-2 “catch” approach to trap IL-2 in close proximity to the cell surface^26^. Wildtype or *Mrtfab*-null OT-I CD8^+^ T cells were coated with IL-2 catch reagent, activated, and surface-proximal IL-2 detected 18h later using fluorescently-labelled IL-2 antibody. Activated OT-I WT cells exhibited large puncta of IL-2 staining, reflecting retention of IL-2 within the cluster; in contrast, the puncta observed in OT-I *Mrtfab*^-/-^ clusters were much smaller (Fig.9C). These data show that MRTF activity potentiates IL-2 signalling by facilitating homotypic cluster formation, allowing effective presentation of IL-2 to recipient cells.

Having demonstrated that aberrant clustering of *Mrtfab*^-/-^ CD8^+^ T cells is associated with defective IL-2 presentation *in vitro*, we evaluated the role of MRTFs in CD8^+^ T cell dynamics *in vivo*. To synchronize T-cell activation as far as possible, we transferred OT-I CD8^+^ T cells into wildtype animals and immunised them by injection with OVA peptides. IL-2Rα expression was comparable in OT-I *Mrtfab*^-/-^ and OT-I WT cells purified from lymph nodes 24h following activation (Fig.S9D). Multiphoton live imaging of popliteal lymph node explants following activation of co-transferred OT-I *Mrtfab*^-/-^ and OT-I WT CD8^+^ T cells labelled with CFSE or SNARF showed that although *Mrtfab*^-/-^ CD8^+^ T cells exhibited similar net displacement over time, their speed was significantly reduced (Fig.S9E). To visualise cell clustering, we generated LifeAct-GFP-expressing OT-I WT and OT-I *Mrtfab*^-/-^ cells, labelled the OT-I WT cells with CellTrace Violet prior to co-transfer, and analysed draining inguinal lymph nodes (Fig.9D). Cells of both genotypes were present in comparable numbers, and following activation by anti-DNGR1-OVA and anti-CD40, many accumulated in clusters of two cells or more. However, the proportion of OT-I *Mrtfab*^-/-^ cells in clusters was significantly reduced (16.9±2.1% versus 31.6±2.8%; Fig.9E), as was the frequency of homotypic interactions between *Mrtfab*^-/-^ cells (17±4.7% cells v 48.54±6.8%)(Fig.9E).

In sum, our data show that MRTF-SRF activity is required for receipt of IL-2 signals by CD8^+^ T cells during the response to LM-OVA infection. This reflects the role of MRTF-SRF signalling in cytoskeletal gene expression, which is required for effective CD8^+^ T cell homotypic clustering *in vivo*.

## DISCUSSION

We have shown that the SRF transcription factor, acting cell-autonomously in partnership with its actin-regulated cofactors, the MRTFs, is essential for the effective response of CD8^+^ T cells to *L. monocytogenes* infection in the mouse. CD8^+^T cells lacking the MRTFs or SRF undergo initial activation and proliferation, but subsequently fail to expand and generate memory cells owing to defective receipt of IL-2 signals. Following activation, *Mrtfab*-null cell generate smaller, less dense clusters with reduced IL-2 retention, and *in vivo Mrtfab*-null CD8^+^ T cells exhibit reduced motility and defective homotypic clustering. Gene expression analysis shows that *Mrtfab*-null cells have substantial deficits in basal cytoskeletal gene expression.

In the LM-OVA infection model, initial TCR activation and proliferation of *Srf*- and *Mrtf*-null CD8^+^ T cells was normal, but expansion was not sustained, and this reflected decreased proliferation rather than increased cell death. Differentiation into effector SLEC and MPEC remained intact, although there was a profound decrease in SLEC expansion resulting in skewing of CD8^+^ effector subsets towards MPEC formation. Related phenotypes have been observed in the LCMV and LM-OVA infection models upon inactivation of IL-2Rα^17–19^, IRF4^22–24, 53^, LFA-1^27–29^, ICAM-1^27, 54^ and PYK2^28^. *Srf*-null CD8^+^ T cells exhibited diminished granzyme B expression, although cytokine expression was not reduced, so it is possible that their cytotoxic function *in vivo* may be compromised. We have not examined this directly, but at least following activation *in vitro*, their ability to induce target cell killing was unaffected. We note that IRF4- and IL-2Rα-deficient CD8^+^ cells are defective in killing *in vivo*^17, 24^, although there are conflicting reports concerning LFA-1^29, 55^. *Srf* is also required for the generation and/or persistence of memory cells, but *Srf-*null CD8^+^ T cells could be generated by inactivation of *Srf* after priming. These cells, which expressed elevated levels of the Tcm markers CD62L, CCR7 and IL-2Rβ also exhibited defective sustained proliferation, indicating that SRF is also required for an effective CD8^+^ T memory cell recall response.

Our findings support a model in which *Srf*- and *Mrtf*-null CD8^+^ T cell phenotypes reflect defective IL-2 signalling. Previous experiments have shown that while the initial proliferative response of CD8^+^ T cells is cytokine-independent^11^, sustained proliferation and SLEC accumulation require IL-2 signalling^17–19^. Even before the decline in proliferation seen in *Srf*- and *Mrtf*-null CD8^+^ T cells, signalling deficits are apparent, including reductions in IL-2Rα and IRF4 expression, reduced activity of the Akt-mTORC1-S6 kinase pathway, and reduced activation of the cytokine effector STAT5. IL-2, like SRF, is required for memory CD8^+^ T cell generation and recall response^14, 56^. Unlike *Il2*-null cells, however, *Srf*-null memory CD8^+^ T cells do not persist following primary infection. Given that *Srf*-null CD8^+^ T cells do generate MPECs, we speculate that SRF may have additional functions required for maintenance of the memory cell population, perhaps involving cell interactions (see below). Like both SRF and IL-2, the transcription factor IRF4 is dispensable for initial CD8^+^ T cell activation and proliferation during infection, but necessary for sustained expansion and SLEC proliferation^22–24^. While initial IRF4 induction is TCR-mediated, *Irf4* is also IL-2 responsive^47^ and it may be that IRF4 production during expansion is also IL-2 dependent.

What is the basis for the defect in IL-2 signalling? *Srf*- and *Mrtfab*-null CD8^+^ T cells produce normal amounts of IL-2 following TCR activation, and IL-2Rα expression is not inherently defective. They retain the ability to respond to excess exogenous IL-2, indicating that the signal transduction apparatus is intact. However, they are unable to respond effectively to endogenously produced IL-2 following activation *in vitro*. Instead, our results suggest that diminished IL-2 signalling in *Mrtfab*-null cells reflects their reduced ability receive endogenous IL-2 signals resulting from defective cell clustering dynamics.

During infection the response to IL-2, including STAT5 phosphorylation and proliferation, is facilitated by cell clustering and LFA-1/ICAM-1 interaction^26, 57^. CD8^+^ T cell clustering is LFA-1/ICAM-1 dependent ^54, 55^ and LFA-1 and its effector kinase PYK2, play an important role in CD8^+^ T cell proliferation and differentiation following LM-OVA or LCMV infection^27–29^. We found that following TCR activation *in vitro*, *Mrtfab*-null CD8^+^ T cells exhibited impaired homotypic clustering, forming smaller and less compact clusters with reduced F-actin content. *Mrtfab*-null CD8^+^ T cell clusters also retain less IL-2. Activated *Mrtfab*-null CD8^+^ T cell dynamics were also impaired in draining lymph node explant cultures: cells moved more slowly, and exhibited a decreased frequency of homotypic interactions. We propose that the LFA-1/ICAM-1-dependent CD8^+^ T cell homotypic interactions required for effective IL-2 presentation *in vivo* in turn depend on the regulation of cytoskeletal dynamics through the MRTF-SRF axis. According to this view, defective IL-2 signalling reflects inefficient IL-2 presentation arising from defective cluster formation. The motility defects in *Mrtfab*-null may cells also contribute to this by decreasing cell encounter frequency, but more work will be required to establish this.

Previous work has shown that paracrine sources of IL-2 are sufficient for effective proliferation in the LCMV model^56^. The cell-autonomous nature of the *Mrtfab*- and *Srf-*null phenotypes suggests that cytoskeletal defects in the recipient cell must be sufficient to impair IL-2 delivery through homotypic CD8^+^ T cell interactions. It therefore remains possible that *Mrtfab* and *Srf* inactivation will also impair delivery of other cytokines, such as IL-12 and interferons, that is mediated by heterotypic interactions^27, 50^.

The immune synapse is also LFA-1/ICAM-1-dependent. However, *Srf* or *Mrtfab* inactivation did not detectably impair initial CD8^+^ T cell activation in the LM-OVA model, indicating that naïve cells are capable of forming a functional synapse. This may reflect other adhesion mechanisms operating in this context. Indeed, following TCR ligation *in vitro*, activated *Srf-* or *Mrtfab-*null CD8^+^ T cells actually exhibited enhanced proliferation, and we speculate that cytoskeletal disruption resulting from MRTF-SRF inactivation might alter the dynamics of TCR signalling complexes in this setting. In the LM-OVA model, however, initial proliferation of *Srf-* or *Mrtfab*-null cells remained normal. Thymocyte positive selection, which also requires TCR signalling, is also unaffected in MRTF-deficient animals (PC, DM and RT, in preparation).

Many studies have established the MRTF-SRF axis as a critical homeostatic regulator of cytoskeletal dynamics, in which the MRTFs respond to state of the actin cytoskeleton by acting as G-actin sensors^31, 33–35^. Gene expression analysis revealed both basal and TCR-induced gene expression of cytoskeletal genes is reduced in *Mrtfab*-null CD8^+^ T cells, as observed in other settings^38, 46^. MRTF inactivation reduced expression of β and γ cytoplasmic actins at both the transcriptional and protein level, having a more pronounced effect on γ-actin. The two cytoplasmic actins are largely functionally equivalent, and it is their differential translational regulation that underpins their specific inactivation phenoptyes^58, 59^. Whether the substantially reduced F-actin content observed in *Mrtfab*-null CD8^+^ T cell clusters reflects the differential effect of MRTF inactivation on β- and γ-actin expression, or more general disruption of cytoskeletal gene expression, will be the topic of future work. We also observed that *Mrtfab*-null CD8^+^ T cells exhibited deficits in their transcriptional response to acute IL-2 stimulation *in vitro*, but their proliferative response to exogenous IL-2 appears unimpaired both *in vitro* and *in vivo*.

We have shown that SRF, acting with its MRTF cofactors, is essential for the effective immune response of CD8^+^ T cells during LM-OVA infection in our mouse model. In humans, inactivation of MKL1/MRTF-A is sufficient to cause immune cell cytoskeletal defects and susceptibility to bacterial infections, but one patient nevertheless successfully resolved a chickenpox infection^60, 61^. The two MRTFs are functionally redundant in many settings^43, 46^, so this observation may reflect residual MKL2/MRTF-B activity in such patients. In conclusion, this study illustrates the importance of cytoskeletal dynamics for the control of cell proliferation in response to IL-2 during the immune response.

## EXPERIMENTAL PROCEDURES

### Mice

C57BL6/J mice were *Elk4*^-/-40, 41^ (SAP-1 null); R26CreERT2 (*tm9(cre/ESR1)Arte*^62^); conditional *Srf*^f/f^ ^63^; conditional *Mrtfa^-/-^Mrtfb^f/f^* ^46^ and transgenic Lifeact-EGFP^64^. *Srf*^f/f^ Tam-Cre (CD45.2), WT Tam-Cre (CD45.1), *Mrtfa^-/-^Mrtfb^f/f^*TamCre (CD45.2), *Elk4*^-/-^ and Lifeact-EGFP animals were crossed to OT-I TCR transgenics (*Tg*(*TcraTcrb*)*1100Mjb*/Crl). OT-I Lifeact-EGFP animals were further crossed to *Mrtfa^-/-^Mrtfb^f/f^* TamCre. For reconstitution, one-week acid-watered *Rag2^-/-^* (RAG2 tm1Fwa) hosts were ^137^Cs-irradiated (2 × 5 Gy) and 24 hours later, bone marrow cells injected into the tail vein. To induce CreER^T2^, mice were fed powdered tamoxifen pellets (Envigo) mixed 4:1 with normal diet to ease absorption for 15 days. Animals were maintained under specific-pathogen–free conditions in The Francis Crick Institute UK Biological Resources Facility. Animal experimentation was carried out under Home Office licences PPL PP0389970, P7C307997, 80/2602 and 70/7982.

### Infection

*L. monocytogenes* expressing ovalbumin (rLM-OVA)^45^ was cultured overnight in brain-heart infusion medium (Fluka 53286-5006) with erythromycin, re-seeded and cultured to OD600 of 0.1-0.2, and diluted in PBS to 10^4^ bacteria/ml for injection. Splenocytes or lymph node cells were enriched for CD8^+^ T cells by negative selection using a CD8 T cell isolation kit (Miltenyi Biotech) based on magnetic cell separation (MACS). 5000 cells (low input) or 2×10^6^ cells (high input) and adoptive transferred by injection into the tail vein. The next day, mice were infected intravenously with 2000 CFU. For recall response, memory CD8^+^ T cells purified by flowsort to 99% purity from splenocytes immunostained with CD8, CD45.1 and CD45.2 were adoptively co-transferred (1:1, 5×10^5^ cells) into naïve syngeneic mice followed by infection with LM-OVA and analysis 6 days later.

### Immunisation

For cell activation and *ex-vivo* analysis of cell clustering, mice were immunised with VacOva (vac-pova, InvivoGen; 50μg/mouse) and LPS (L6529 Sigma; 30μg) co-injected subcutaneously into the hock; and for live microscopy, with anti-DNGR1 397-OVA fusion protein^65^ (2μg; gift from Sandra Diebold) and anti-CD40 (1C10; 25μg) co-injected into the footpad.

### Flow cytometry

Tissues were disaggregated through a 70μm nylon mesh in cold RPMI 10% FCS. Blood was collected into Sarstedt EDTA KE/1.3 tubes before red blood cell lysis. Cells were stained in ice-cold FACS buffer (1% FCS, 2mM EDTA in PBS) with combinations of fluorochrome-conjugated antibodies and analysed using BD-Fortessa Instruments and FlowJo 9.9 software. For intracellular staining, cells were fixed and permeabilised using eBioscience intracellular fixation and permeabilisation kit (BD Biosciences). For pSTAT5 and pERK staining, cells were fixed in 2% paraformaldehyde, washed and permeabilised in ice-cold methanol for 30 min, washed twice in PBS, 10% FCS and stained for 1 h. Samples were run on LSR-IIB or Fortessa II (BD Biosciences) and analysed with FlowJo software (BD Biosciences).

### Cell isolation

Where indicated, T-cells were sorted to >97% purity as judged by cell surface marker expression on a BD INFLUX or FACS ARIA-III using BD FACS Diva software for subsequent analysis. Coulter CC Size standard beads (Beckman Coulter) were used for calculating cell numbers. For clustering experiments, CD8^+^ naïve T cells were enriched from lymph node or spleen of bone marrow reconstituted mice by MACS.

### Cell stimulation

Cells were cultured in R10 medium, (RPMI 1640 containing 10% heat inactivated FCS, 50μM 2-mercaptoethanol with penicillin and streptomycin, (Sigma-Aldrich) and incubated at 37°C, 10% CO2. For cell proliferation assays, purified naïve CD8^+^ T cells were labelled with CFSE (5μM, Molecular Probe) for 15 min in RPMI 1640 without serum and washed in R10 medium to stop the reaction. Cells were cultured on plate-bound anti-CD3 (2C11) and anti-CD28 (37.51) (5μg/ml) or stimulated with OVA peptides of decreasing affinity (N4 > Q4 > V4) for the OT-I TCR: N4 (SIINFEKL), Q4 (SIIQFEKL) and V4 (SIIVFEKL)^66^.

For intracellular cytokine staining, cells were stimulated with 10nM SIINFEKL (OVA) peptide for 5h at 37°C in the presence of Brefeldin A (Sigma–Aldrich) for the last 2h before staining. Release of IL-2 was measured by IL-2 ELISA (M2000 R&D).

For *ex-vivo in vitro* culture, splenocytes were harvested from 3-day post-infection mice and cultured for a further 72h at 1-2×10^6^ cells/ml in media alone, or supplemented with 20ng/ ml rmIL-2 or 20 ng/ml rmIL-12 before analysis by Flow cytometry.

### *In vivo* IL-2 modulation

IL-2/mAb complexes were generated by incubating murine rIL-2 (rmIL-2, Immunotools) with S4B6 anti-IL-2 monoclonal antibody at a 2:1 molar ratio (1.5μg/ml IL-2, 50μg/ml S4B6) for 15 minutes at room temperature. IL-2/S4B6 complexes or IgG2α control antibodies (both eBioscience) were injected i.p.

### *In vivo* T-T clustering assay

Purified OT-I Lifeact-EGFP CD8^+^ T cells were labelled with 5 μM of CellTrace Violet for 30 min at 37°C and then transferred into recipient mice by tail vein injection. Mice were immunised a day later and the draining lymph nodes were harvested 24 hours later. Lymph nodes were in fixed in 4%PFA overnight at 4°C and then transferred to a solution of 30% sucrose 1x PBS at 4°C overnight. Lymph nodes were then embedded and snap frozen in OCT (VWR 361603E) and stored at −80°C. 30 μM tissue sections were produced and mounted into ProLong Diamond Antifade (Thermofisher). Sections were scanned using a Zeiss LSM880 inverted with an airyscan module. (alpha Plan-apochromat 63x /1.46 Oil Korr M27 objective lens, 0.5 μM intervals, x,y: 0.033 μm). Distance between cells and number of direct connections between cells were measured using a custom Cellprofiler pipeline. Briefly, multichannel confocal images were split into single-channel images and processed for background removal in FIJI. Cells were manually segmented using the FIJI Labkit plugin using the GFP channel. The output of the Labkit segmentation produced an image in which each segmented cell had a unique integer pixel value. Segmentation and individual channel images were then imported in CellProfiler v4.2.4, the segmented cells were converted into ‘Objects’ using the ConvertImageToObject module. The fluorescence intensity of the Violet channel was measured inside each object using the MeasureObjectIntensity module and used to filter the objects into the Violet positive and Violet negative categories using the FilterObject module. The distance between cells and the number of neighbouring cells within a 10pixel radius was then measured using the MeasureObjectNeighbors module.

### Immunohistochemistry

For staining of spleen sections, OCT embedded fixed spleens injected with either OT-I WT or OT-I *Mrtfab*^-/-^ T cells labelled with cell tracker deep red dye were prepared as above. 8μm frozen sections were air dried for 30 min at room temperature in the dark and rehydrated for 3 min in PBS. Sections were blocked and permeabilised in blocking buffer 0.3% Triton X-100, 3% BSA in PBS for 1 hour at room temperature followed by incubation with primary antibody diluted in blocking buffer overnight at 4°C. After 3x washesin 1x PBS, sections were incubated for 1 hour with secondary antibody followed by 3 washes in PBS and mounting with ProLong Diamond Antifade.

### Multiphoton live microscopy of explanted popliteal lymph nodes

For analysis of cell migration *in vivo*, *Mrtfab*^-/-^ and WT OT-I cells were labelled with 2μm CFSE (C34554 Invitrogen) or 6μM SNARF (S22801 Invitrogen) and 2×10^6^ cells mixed in 1:1 ratio were transferred into CD45.1/CD45.2 recipients. One day following immunisation, popliteal draining lymph nodes were removed and immobilized on coverslips with hilum facing away from the objective. Lymph nodes were continuously perfused with warmed (37°C) Roswell Park Memorial Institute (RPMI) 1640 medium without Phenol Red (GIBCO, Invitrogen) bubbled with carbogen (95% O2 5% CO2). Images of live cells were taken using a LSM Zeiss 710 multiphoton, 20x NA 1.05 water immersion objective and a pulsed Ti:sapphire laser (Spectra Physics MaiTai HP DeepSee) voxel size:1.6605×1.6605×3μm^3^, to a depth of about 100 μm from the capsule.Tracking of live cells was analysed using Imaris 9.5 using the spot tool applying gaussian filter (2μm) and background subtraction (6μm). Cells that could be detected continuously for 15 min were analysed and the average speed and total displacement were measured.

### *In vitro* T-T clustering assays

Antibody-induced clusters were generated by culturing MACS-purified naïve OT-I T-cells on plate-bound anti-CD3 and anti-CD28 both at (5μg/ml) for 24hrs in the presence or absence of anti-IL-2 blocking antibody (JES6-1A12) (10μg/ml), before transfer of cells and supernatant to non-coated wells with or without anti-CD11a/LFA-1 (M17.4)(10μg/ml) blocking antibody and ^r^mIL2 (20ng/ml) for the times indicated. Pharmacologically-induced clusters were generated by culturing T-cells in 25 ng/ml Phorbol 12,13-dibutyrate (PDBu) (P1269 Sigma) and 500ng/ml ionomycin (I0634 Sigma) for 18h. Clusters were then cultured for an additional 5 hours with or without anti-IL-2 or anti-LFA-1 blocking antibodies or with mIL-2 (20ng/ml) for the last 15 min or with Pyk2/FAK inhibitor PF431496 (5μM) for the last 1h. Following incubation with anti-IL2 blocking antibody some cells were restimulated with ^r^hIL-2 (Sigma) for 15 min. Clusters were visualised using ZEISS Observer D1 AxioCam. For fluorescence staining, activated T cells were transferred to poly-L-lysine-coated Matek dishes (50μg/ml), incubated 15 min at 37C with 5% CO2 and fixed with 2% PFA for 10 min. Clusters were washed in PBS, blocked in 3% BSA, permeabilised in 0.2%Triton and then immuno-stained with anti-pericentrin and counterstained with Texas Red phalloidin and DAPI. For the IL-2 catch assay, purified CD8^+^ T-cells were coated with mouse IL-2 catch reagent from a mouse IL-2 secretion assay detection kit (Miltenyi Biotec) by incubating T cells in a 20x dilution of IL-2 catch reagent in R10 medium for 15 min on ice, washing once in R10 medium and then stimulating with PDBu/Iono to form T-cell clusters as above. After fixation, cells were stained with phycoerythrin-conjugated IL-2 detection antibody (1:20 dilution in R10 medium).

Stained clusters were mounted in Mowiol with coverslips and analysed by confocal microscopy (Zeiss LSM710 inverted). Single-plane bright-field images and Z stacks of the appropriate channel of fluorescent images (0.37μm intervals) were acquired with Plan Apochromat 40x/1.3 oil Ph3 M27 Objective lens. Acquired images were analysed using Imaris 9.6. Images were processed using Imaris background subtraction filter for each channel. Additionally, a gaussian and median filter was applied on the channel of interest. For *in vitro* generated cluster, cells were detected using the Imaris spot tool on the DAPI channel, distance between cells was measured using the spots positions. The mean intensity of the phalloidin staining and the cell sphericity were measured for each individual cell using the Imaris cell detection tool. For the IL-2 catch analysis a median filter was first applied on the images.The Imaris surface tool was used with two volume thresholds (5.04 and 10.1 μm^3^) to create three colour-coded classes of volumes of IL-2 signals in each cluster.

### Adhesion assay

Binding of ICAM-1 complexes to MACS purified OT-I CD8^+^ T cells. Binding of soluble ICAM1-Fc-F(ab’)2 complexes were generated by diluting APC-labeled goat anti-human IgG F(ab’)2 fragments (109-135-098, Jackson Immunoresearch) 1:6.25 with ^r^mICAM1-Fc (200µg/ml final) in HBSS and incubated for 30 min in HBSS at 4°C. Splenocytes were rested for 3 h in IMDM, 5% FCS at 37°C, centrifuged and resuspended in HBSS, 0.5% BSA. Each adhesion reaction (50 µl) contained 20×10^6^ cells/ml, 25 µg/ml ICAM-1 complex and the appropriate stimulus and was incubated at 37°C for the indicated times. OT-I CD8^+^ T cells were stimulated with anti-CD3 (10 μg/ml)). Cells were fixed in PFA for 20 min and binding of ICAM-1 complexes to T cells analyzed by flow cytometry.

### Cytolytic assay

For CD8^+^ T effector differentiation, purified naïve OT-I CD8^+^ cells were activated on plate-bound antiCD3/CD28 (10μg/ml) for 48h, washed three times, then cultured with 20ng/ml recombinant IL-2 for an additional 5 days (rmIL-2, Immunotools). CD8 effector cells were mixed with CFSE-stained EL4 target cells, pulsed or not with 1μg/ml of SIINFEKL peptide, and incubated for 3h before fixation and staining for intracellular active caspase-3 in EL4 target cells.

### BrdU incorporation

Mice were injected (i.p) either at day 1 or day 6 post-infection with 1mg of BrdU (BD Biosciences); 48h later, splenocytes were stained for surface markers and BrdU incorporation according to the manufacturer’s protocol.

### RT-qPCR

Total RNA was isolated using the GenElute kit (Sigma) and treated with DNA’ase I. cDNA was synthesized and quantitative PCR performed using SYBR Green Express and a QuantStudio 5 qPCR instrument (Life Technologies). Primers were:

*Actb* intronic: F-CGTAGCGTCTGGTTCCCAAT;
R--GTGTGGGCATTTGATGAGCC
*Actb* exonic: F- CGCCACCAGTTCGCCAT;
R- CTTTGCACATGCCGGAGC
*Actg* intronic: F- CTGGCCGAGGACATTTTCTG;
R- GAAGAAGCCCCGGAATTAGC
*Actg* exonic: F- ATGGAAGAAGAAATCGCCGC;
R--AGGGTCAGGATACCCCTCTT
*RpS16*: F- CGCACGCTGCAGTACAAGTTACT;
R--ACATGTCCACCACCCTTCACAC
*IL-2* exonic F- TCAGTGCCTAGAAGATGAACTTG;
R- TCAAATCCAGAACATGCCGC
*Srf* exonic F- CACCTACCAGGTGTCGGAAT;
`R-` GCTGTGTGGATTGTGGAGGT

### RNAseq and bioinformatics

OT-I WT and OT-I *Mrtfab*^-/-^ were sorted from live CD8^+^ gated lymph nodes cells derived from tamoxifen-fed bone-marrow reconstituted mice. Cells were activated with plate-bound anti-CD3/CD28 (5μg/ml) for 24h, and washed “IL-2 quiesced” and resuspended overnight in media supplemented with IL-12 cytokine (20ng/ml). The next day cells were activated with rIL-2 (20ng/ml) for the times indicated followed by RNA extraction. For RNAseq, RNA was prepared using total RNA Sigma GeneElute columns (Sigma). Libraries were prepared using Nugen cDNA synthesis kit.

FASTQ files for each sample were processed via the nf-core/rnaseq pipeline (v3.5)^67^ against the Mus Musculus mm10 genome build and RefSeq annotation GCF_000001635.26_GRCm38.p6 using STAR with strandedness set to unstranded. BigWig coverage files were generated using deeptools v3.0.0 with normalisation factors calculated as described^68^.

Per-gene read counts were retrieved using the R/Bioconductor (v4.0.3) library GenomicAligment (v1.26.0) and the function summarizeOverlaps^69^ against the GCF_000001635.26_GRCm38.p6 RefSeq annotation.

All bioinformatics analysis methods and code are available at GitHub repository link https://github.com/fgualdr/maurice_etal_2023_MRTF_SRF_IL2delivery/

Total and intronic read counts per gene were collected using the summarizeOverlaps method and used for downstream analysis. Sample normalization was achieved by selecting invariant genes across samples/conditions^37^ using the R library GeneralNormalizer. Differentially regulated genes were selected using DESeq2 (R/Bioconductor package version 1.26.0; R version 3.6.2) after turning off the default normalization that DESeq2 applies. Genes differentially expressed considering either all or intronic read counts with an associated absolute Log2FoldChange greater than 1 and an adjusted p-value less than or equal to 0.05 were collected and clustered according to the z-score across conditions (refer to individual plots and figures). Unsupervised clustering was achieved using dimensionality reduction coupled with Louvain community detection. Gene set enrichment analysis was conducted against the MSigDB (msigdb_v7.5.1) computing the hypergeometric enrichment and the Standardized Jaccard index.

Raw and processed RNAseq data are in GEO, accession GSE241689.

### Statistical Analysis

Data were analysed with Graph Pad Prism 6. Bar and dot charts are expressed as mean ± SEM and data analysed using the unpaired and paired parametric *t* test. Statistical significance is indicated on the Figures: *, p <0.05; **, p<0.01; ***, p<0.001; ****, p<0.0001; ns, not significant.

### Antibodies and Cytokines

Antibodies were from:

eBioscience: CD8 (53-6.7), CD122 (5H4),TCRβ (H57-597), IRF4 (3E4), IFNγ (XMG1.2), CD44 (IM7), CD122 (5H4), CD5(53-7.3) CD62L (MEL-14), CXCR3 (173), CCR7 (4B12), NKG2D (CX5), TNF-α (MP6-XT22), GranzymeB (NGZB), IRF4(3E4).

BD biosciences: CD11b (M1/70), CD45.1 (A20), CD45.2 (104), Bcl-2 (3F11), CD4 (RM4-5), CD127 (SB/199), KLRG1 (2F1), αβTCR (β-chain. H57-597), CD45.2 (104), Caspase-3 (C92-605), HSA (M1/69), IL-2 (JES6-5H4); pan ERK(16), CD25 (PC61.5), CD132 (TUGm2),CD3ε (2C11),CD24 (M1/69), CD25 (7D4), CD69 (H1.2F3), phosphoSTAT5 (pY694) (47), panSTAT5 (89), KLRG1 (2F1), TNFα (MP6-XT22), CD28(37.51).

Biolegend: CD11a/CD18 (H155-78), CD62L(MEL-14), CD69(H1.2F3), Streptavidin Brilliant violet 421, CCL5 (2E9).

Cell Signaling: pAKT(T308)(D25E6), pS6 240/244 (D68F8),pS6 235/236 (D57.2.2E), p44/42-ERK (D13.14.4E), pStat5 (pY694),Stat5(D206Y)

APC-labeled goat anti-human IgG F(ab’)2 fragments (109-135-098,Jackson Immunoresearch), rmICAM1-Fc (R&D systems).

**Blocking antibodies**: IL-2 blocking (JES6-1A12), IL-2 monoclonal antibody (S4B6), LFA-1/CD11a blocking (M17/4)

**Western blot antibodies** anti-SRF (G20, Santa-Cruz Biotechnology), anti-panERK (BD biosciences) anti-MRTF-B (Bethyl A302-786A,a-MKL2), anti-γ actin (cytoplasmic, lot1108,β-CYA,gift from Christine Chaponnier and Michael Way), anti-β actin (Sigma A2228)

**Cytokines**: Cytokines: rmIL-2 (12340026 ImmunoTools), rhIL-2 (Proleukin; Chiron B.V., Leiden, The Netherlands), rmIL-12 (419-ML-050 R&D)

## Supporting information

Supplementary Table S1. Gene expression in CD8+ T cells.

Supplementary Table S2. Gene cluster analysis

## ACKNOWLEDGEMENTS

We thank Adrian Hayday, Caetano Reis e Sousa, Gitta Stockinger, Carola Vinuesa, and lab members for helpful discussions and suggestions during the project and/or comments on the manuscript. We thank Santiago Zelenay for advice, reagents and assistance with the Listeria experiments, and Sandra Diebold for the anti-DNGR1 397-OVA fusion protein. We are grateful to the following Crick Science technology platforms and staff: Phil East and Aengus Stewart at Bioinfomatics and Biostatistics for processing raw RNAseq data; Jimena Perez-Lloret, Jerome Nicod and Advanced Sequencing for library preparation and RNAseq; Sukhveer Purewal and Derek Davies at Flow Cytometry; Camille Charoy at Advanced Light Microscopy for fluorescent image acquisition and analysis; and Clare Watkins and Julie Bee and the Biological Resources Facility for expert support. This work was supported by the Francis Crick Institute which receives its core funding from Cancer Research UK (CC2102), the UK Medical Research Council (CC2102), and the Wellcome Trust (CC2102). This research was funded in whole, or in part, by the Wellcome Trust CC2102. For the purpose of Open Access, the author has applied a CC BY public copyright licence to any Author Accepted Manuscript version arising from this submission.·The authors have no conflicts of interest.

## AUTHOR CONTRIBUTIONS

DM constructed the conditional SRF and MRTF mouse strains; conceived and designed the cotransfer-infection protocol and designed, carried out and interpreted infection, signalling, and imaging experiments; PC designed, performed and interpreted experiments; FG designed and conducted the bioinformatic analyses; BF set up the multiphoton life microscopy system and conceived and performed the *in vivo* imaging. RT conceived the project, designed and interpreted experiments, and wrote the paper with DM.

## SUPPLEMENTARY FIGURE LEGENDS

**Figure S1.**
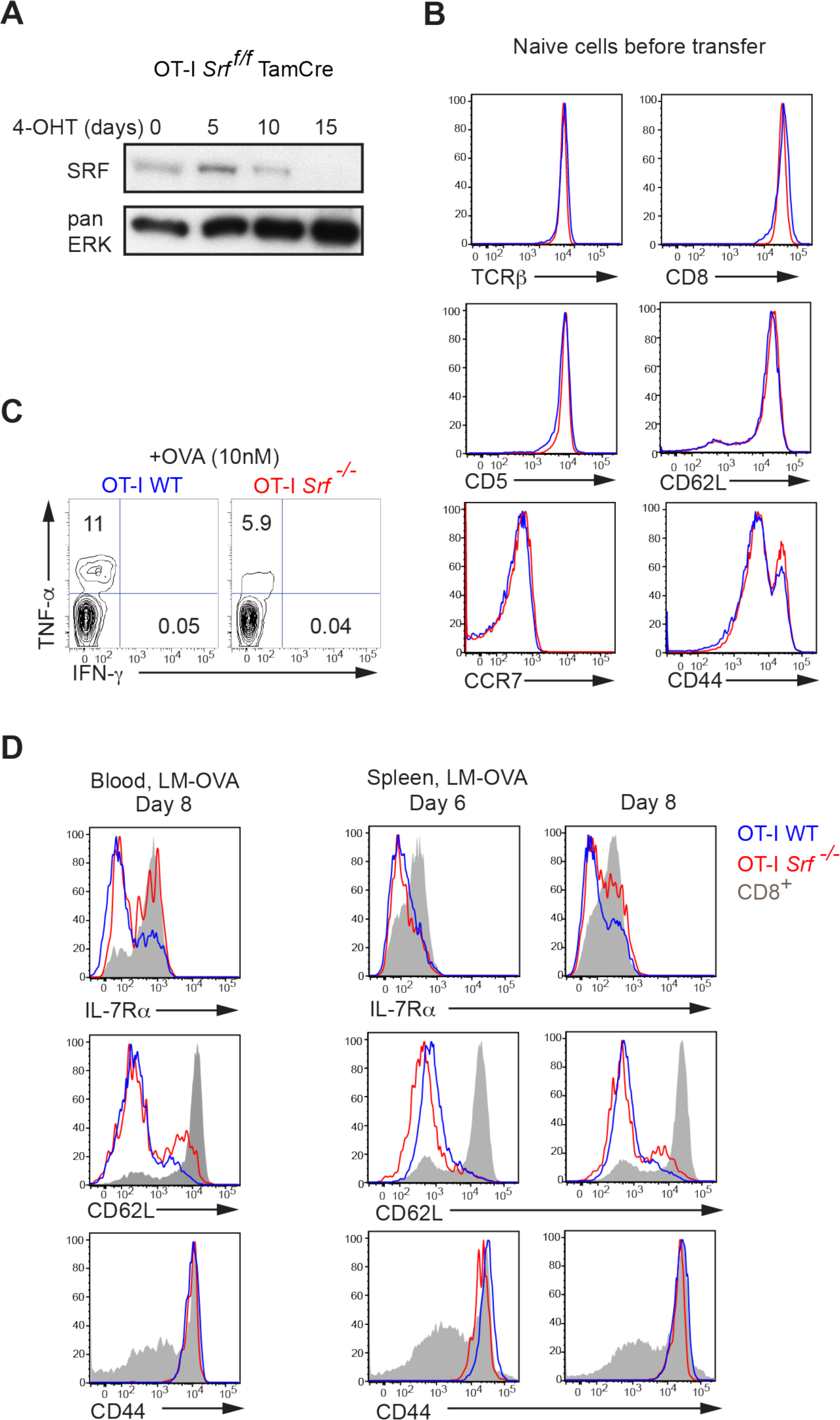
Phenotypic analysis of OT-I *Srf*^-/-^ CD8^+^ T cells. **(A)** Time course (days) of SRF expression in tamoxifen-fed animals. **(B)** OT-I WT and OT-I *Srf^-/-^* CD8^+^ T cells were MACS-purified from the spleens of tamoxifen-treated bone marrow reconstituted mice and stained for TCRβ, CD8, CD5, CCR7, CD62L and CD44 expression. **(C)** Purified OT-I cells were stimulated for 5h with SIINFEKL OVA peptide and analysed for intracellular IFN-γ and TNF-α expression. **(D)** IL-7Rα, CD62L and CD44 expression *in vivo* after LM-OVA infection in spleen and blood.

**Figure S2.**
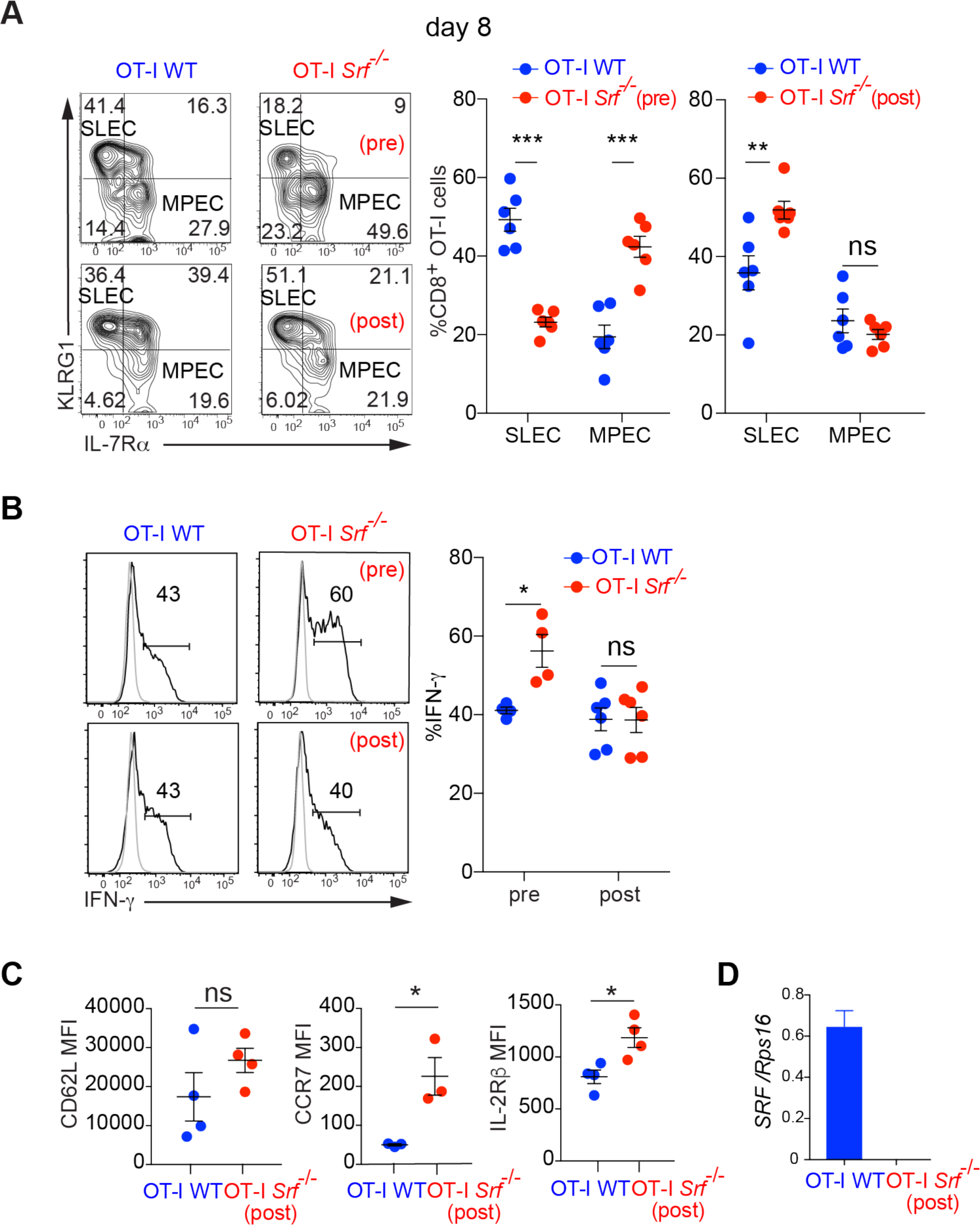
*Srf* is required for effective SLEC accumulation. **(A)** MPEC and SLEC generation in infected splenocytes. Wildtype cells were co-transferred with cells in which *Srf* was inactivated either prior to transfer (OT-I *Srf*^-/-^ (pre)) or after transfer (OT-I *Srf*^-/-^(post)), and infected as outlined in Fig.1A (OT-I *Srf*^-/-^(pre)), or Fig. 2A (OT-I *Srf*^-/-^(post)). Data show mean values ± SEM; data points represent individual mice; n=6. **(B)** Cytokine production by infected splenocytes stimulated with SIINFEKL peptide (OVA; 10nM) for 5 hours. Wildtype cells were co-transferred with cells in which *Srf* was inactivated either prior to transfer (OT-I *Srf*^-/-^(pre)) or following transfer and infection (OT-I *Srf*^-/-^(post)). Data show mean values ± SEM; data points represent individual mice. **(C)** Memory-like OT-I WT and *Srf*^-/-^(post) cells, defined by CD62L, CCR7 and IL-2Rβ expression. **(D)** *Srf* mRNA levels in sorted OT-I splenocytes at day 50 post-infection.

**Figure S3.**
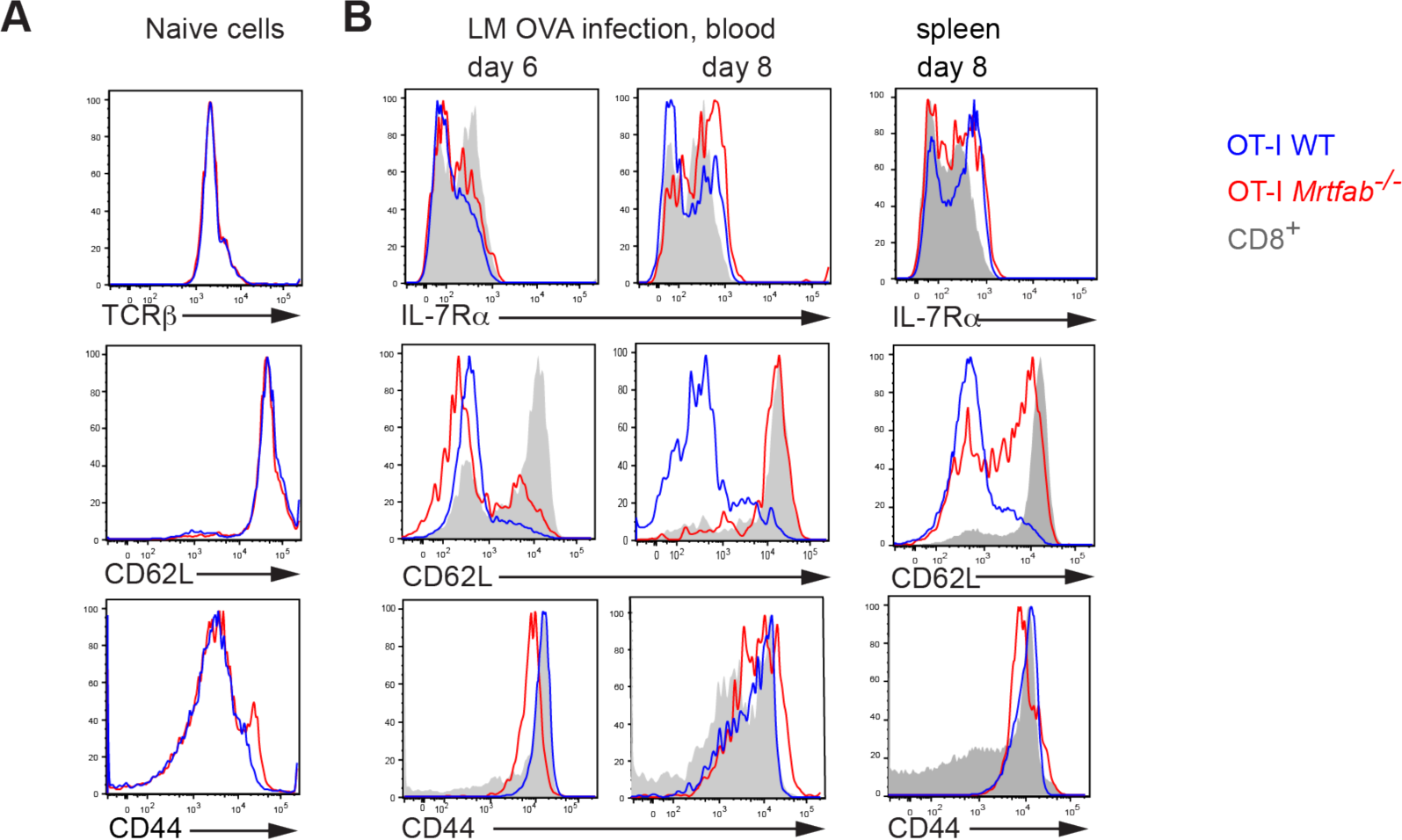
Phenotypic analysis of naïve OT-I *Mrtfab*^-/-^ CD8^+^ T cells. **(A)** OT-I WT or OT-I *Mrtfab^-^*^/-^ cells were MACS-purified from the spleens of tamoxifen-treated bone marrow reconstituted mice and stained for TCRβ, CD62L and CD44 expression. **(B)** Expression of IL-7Rα, CD62L and CD44 on OT-I WT or OT-I *Mrtfab^-^*^/-^ CD8^+^ T cells following LM-OVA infection in blood and spleen.

**Figure S4.**
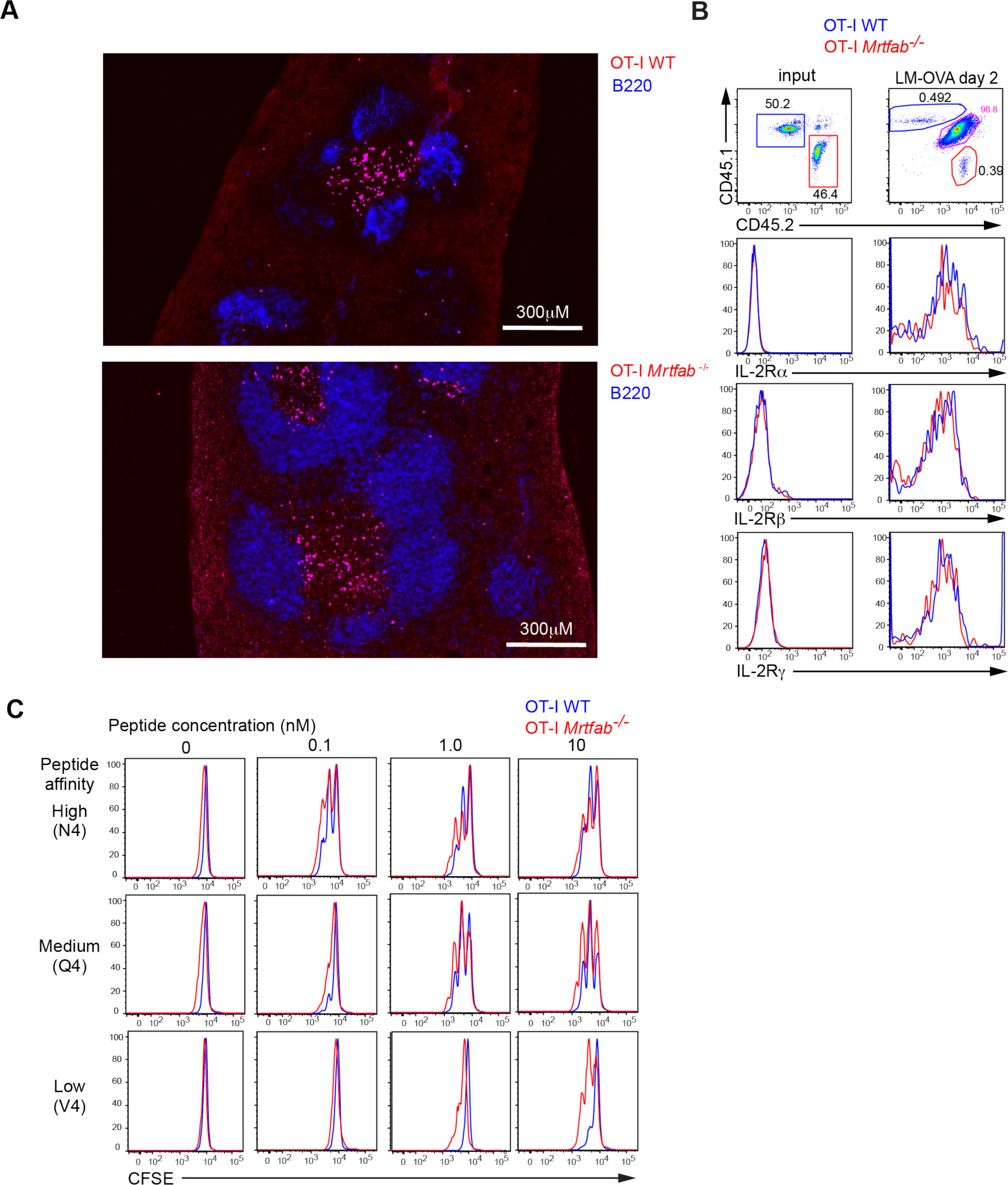
OT-I *Mrtfab*^-/-^ cells localisation after LM-OVA infection. **(A)** OT-I *Mrtfab^-^*^/-^ and OT-I WT CD8^+^ T cells both localise to the periarteriolar lymphoid sheath. OCT-embedded sections of spleens from mice injected with OT-I *Mrtfab^-^*^/-^ and OT-I WT CD8^+^ T cells labelled with cell-tracker deep red harvested 3d following LM-OVA infection, and counterstained for B220 to reveal B cell follicles. Scale bar 300μm. **(B)** Expression of IL-2Rα, β and γ chains in OT-I WT and OT-I *Mrtfab*^-/-^ cells in cells before and 2 days after LM-OVA infection. **(C)** Representative CFSE division profiles of MACS-purified OT-I WT or OT-I *Mrtfab^-^*^/-^ cells, activated for 48h by indicated concentrations of OVA peptides of decreasing affinity (N4 > Q4 > V4) for the OT-I TCR.

**Figure S5.**
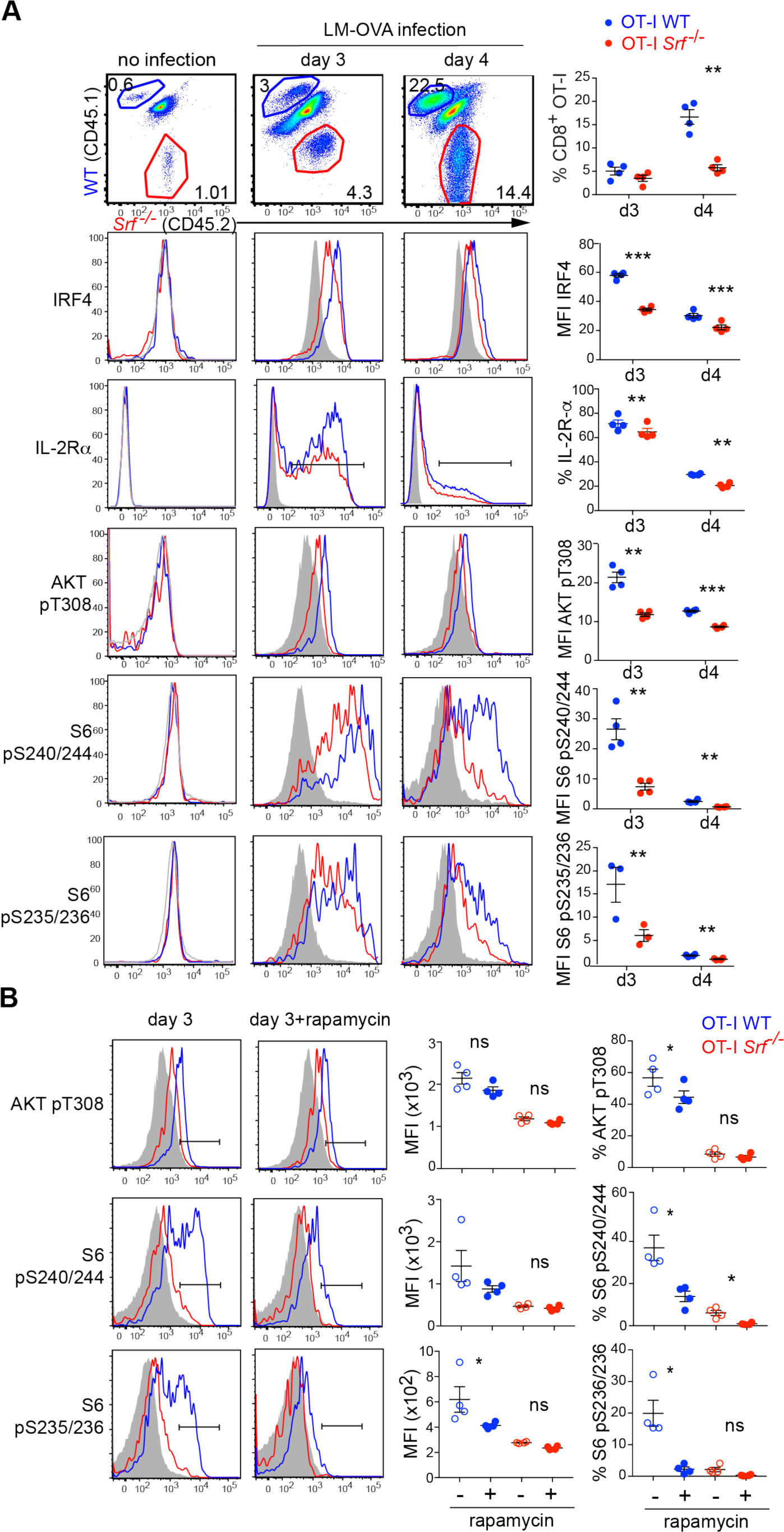
IL-2 signalling is defective in OT-I Srf^-/-^ during infection. **(A)** Quantification of protein expression in OT-I CD8 splenocytes populations from mice co-transferred with OT-I WTCD45.1 and OT-I *Srf*^--/-^ CD45.2 (1.10^6^ cells) and harvested at day3 or day4 p.i, presented as in Fig.5. Each data point represents a single mouse (n=4) in one representative experiment. Data show mean values ± SEM; Statistical significance, paired t test. Experiment has been done ≥3 times. **(B)** Expression of pAKT, pS6240/244 and 235/236 in OT-I WT (blue) and OT-I *Srf^-/-^* (red) and endogenous CD8 T (grey-filled) cells in mice infected with LM-OVA. Splenocytes harvested 3 days after infection are treated or not with rapamycin (20nM) for 20 min ex vivo followed by antibody staining and analysis by flow cytometry. Statistical significance, paired t test.

**Figure S6.**
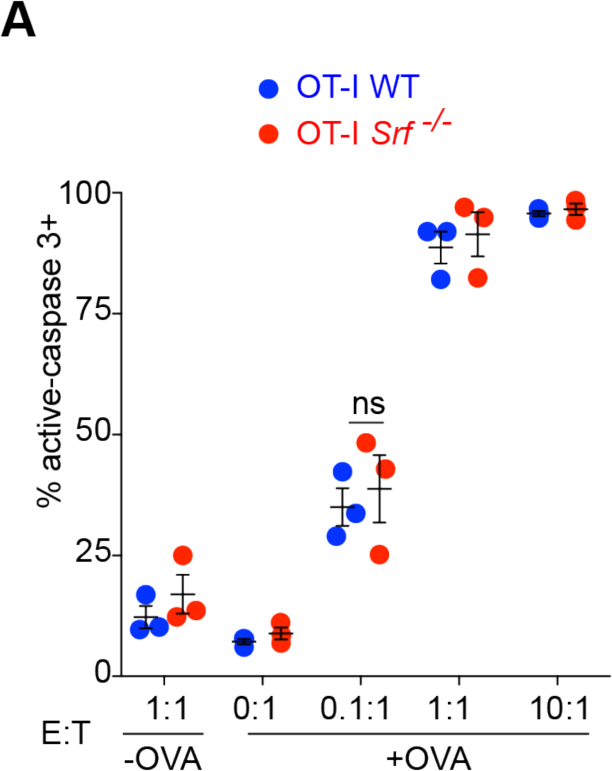
*In vitro* cytotoxic assay. **(A)** *In-vitro* killing by OT-I WT and OT-I *Srf*^-/-^ effector CD8^+^ T cells activated by plate-bound anti-CD3/CD28 and cultured for 7 days in IL-2 (20ng/ml), then incubated with OVA-pulsed target EL4 cells at different effector:target cell ratios. Killing of EL4 cells measured by intracellular caspase-3 staining. Data show mean values ± SEM; data points represent individual mice.

**Figure S7.**
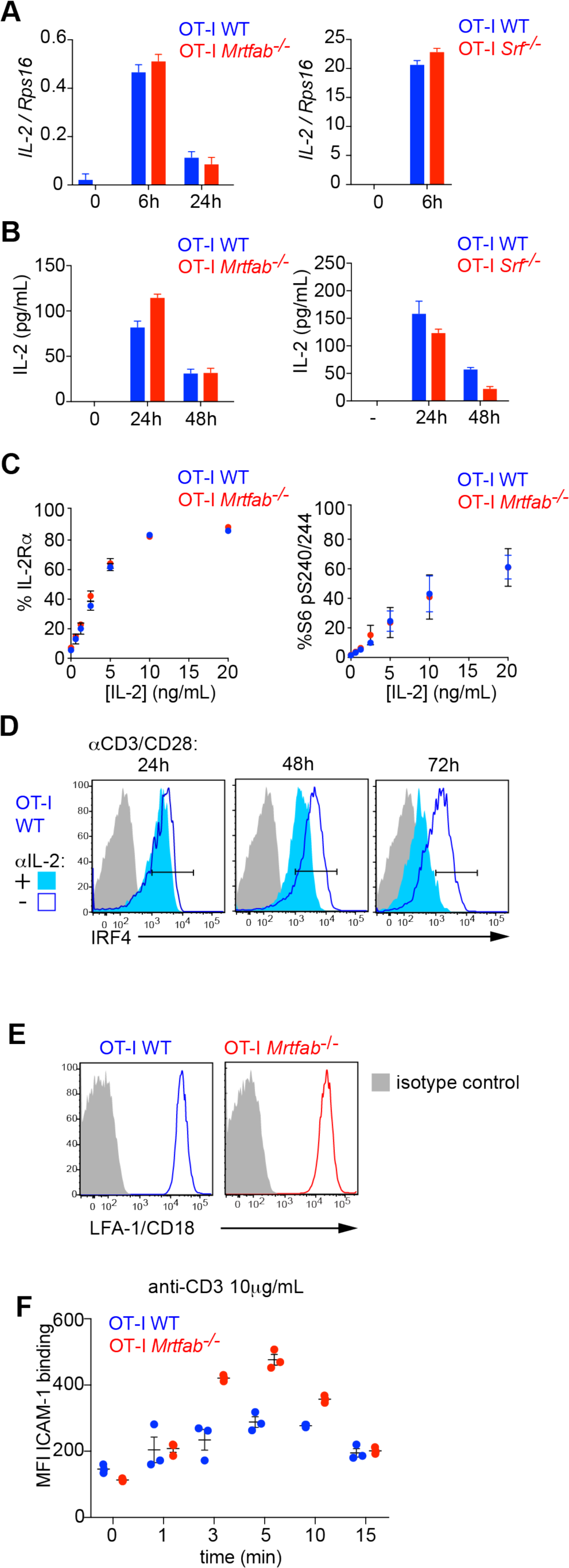
IL-2 cytokine production and signalling is intact in activated OT-I *Srf*^-/-^ and OT-I *Mrtfab*^-/-^ cells. **(A)** Plate-bound anti-CD3/CD28 TCR-induced transcription of IL-2 mRNA in sorted OT-I lymph nodes cells. **(B)** IL-2 production measured by ELISA in sorted CD8^+^ T lymph nodes cells upon plate-bound anti-CD3/anti-CD28 activation. Experiment done twice, each bar represents triplicate for one representative experiment +/-SEM. **(C)** Expression of IL-2Rα (CD25) and pS6240/244 in sorted OT-I lymph node cells preactivated for 24h with plate-bound anti-CD3/CD28, washed and replated with a titration of IL-2 cytokine for 24h. **(D)** Expression of IRF4 in sorted OT-I lymph node cells preactivated for 24h with plate-bound anti-CD3/CD28 in the presence or absence of anti-IL-2 blocking antibody. **(E)** Expression of LFA-1 in TCR-activated OT-I cells. **(F)** Time course of ICAM-1 binding to TCR-activated OT-I cells

**Figure.S8.**
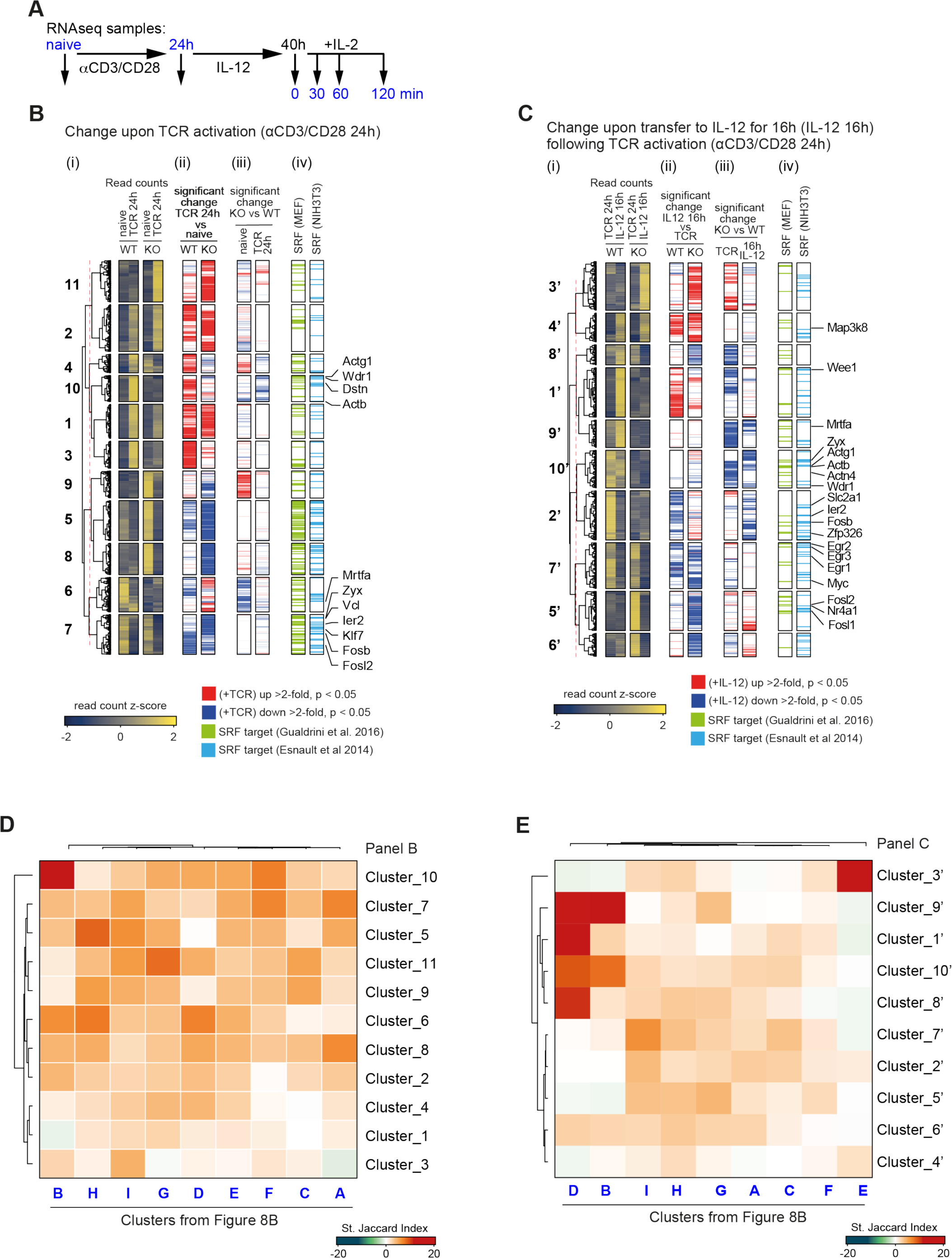
Gene expression deficits in OT-I *Mrtfab^-/-^* cells. **(A)** Purified naïve wildtype or *Mrtfab*-null OT-I CD8^+^ T cells were stimulated as shown before RNAseq analysis^52^. **(B)** (i) Normalized z-scored read counts of genes differentially expressed in WT or *Mrtfab*-null cells following TCR activation for 24h, grouped by unsupervised clustering (clusters 1-11). Data show mean values ± SEM of three biological replicates. (ii) Genes showing significant changes upon TCR activation in wildtype (left) and *Mrtfab*-null cells (right). (iii) Genes whose expression is impaired by MRTF inactivation in naïve cells (left) or TCR-activated cells (right). (iv) Genes identified as candidate SRF targets in TPA-stimulated MEFs^37^ or serum-stimulated NIH3T3 fibroblasts^36^. **(C)** (i) Normalized z-scored read counts of genes differentially expressed in WT or *Mrtfab*-null cells upon resting in IL-12 following TCR activation (clusters 1’-10’). Data show mean values ± SEM of three biological replicates. (ii) Genes showing significant changes following culture in IL-12 in wildtype (left) and *Mrtfab*-null cells (right). (iii) Genes whose expression is impaired by MRTF inactivation, in TCR-activated cells (left) or TCR-activated/rested cells (right). (iv) Genes identified as candidate SRF targets in TPA-stimulated MEFs^37^ or serum-stimulated NIH3T3 fibroblasts^36^. **(D)** Overlap testing analysis, displayed as standardised Jaccard score, of the relation between gene groups differentially expressed upon TCR-activation of WT or *Mrtfab*-null cells (clusters 1-11 in panel B) and those differentially expressed between WT and *Mrtfab*-null TCR-activated/rested cells with or without stimulation by IL-2 (Fig.8B, groups A-I). **(E)** Overlap testing analysis, displayed as standardised Jaccard score, of the relation between gene groups differentially expressed in WT or *Mrtfab*-null cells upon resting in IL-12 following TCR activation (clusters 1’-10’ in panel C) and those differentially expressed between WT and *Mrtfab*-null TCR-activated/rested cells (Fig.8B, groups A-I).

**Figure S9.**
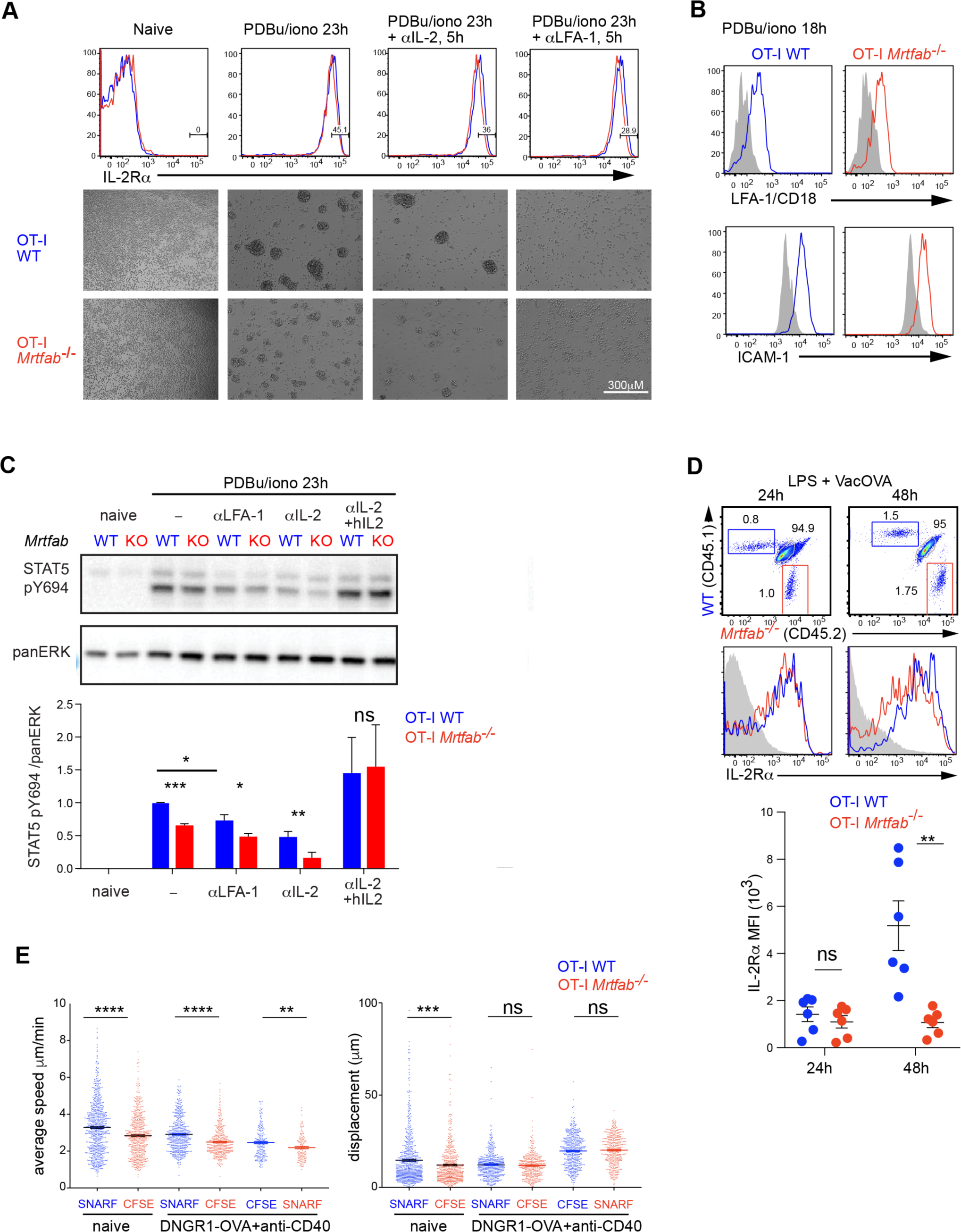
*Mrtfab*-null CD8^+^ cells exhibit defective homotypic clustering *in vitro* and defective motility *in vivo*. **(A)** MACS purified OT-I CD8T splenocytes were stimulated with PDBu/Iono to induce cluster formation. At 18hr, cells were treated or not with anti-LFA-1 (,10μg/ml) or anti-IL-2 blocking antibody (JES6-1A12,10μg/ml) for a further 5h. Top, IL-2Rα expression assessed by flow cytometry. Bottom, cluster formation at 23h with αIL-2 or αLFA1 added for the last 5h. **(B)** LFA-1 and ICAM expression in cells stimulated as in (A) **(C)** STAT5 pY694 expression in clusters induced as in (A) or similarly treated at 18h with anti-IL-2 blocking antibody for 5h with or without human IL-2 (20ng/ml) for 15 min. Bottom, quantitation of 3 independent experiments +/- SEM. **(D)** Mice adoptively co-transferred with CD45.1 OT-I WT and CD45.2 OT-I *Mrtfab*^-/-^ CD8^+^ T cells were immunised one day later, and inguinal lymph nodes were analysed at 24h and 48h later. IL-2Rα expression in endogenous control (grey), OT-I WT (blue) and OT-I *Mrtfab*^-/-^ (red) gated populations is shown, with MFI quantified as mean values ± SEM;, paired t test as above. Each data point represents an individual mouse. **(E)** Speed of OT-I *Mrtfab*^-/-^ T cells is decreased in lymph-node by comparison to OT-I WT cells. Wild-type mice were co-injected with a combination of CFSE- or SNARF-labelled OT-I WT and OT-I *Mrtfab*^-/-^ T cells and immunised with anti-DNGR1-OVA plus anti-CD40 or PBS vehicle. Popliteal lymph nodes were removed 24h later and imaged for 30 min time lapses. Dots represent the average speed or track displacement of individual cells tracked over 15 min using Imaris software. Data shows 3 independent experiments. Similar results were obtained when the dyes were exchanged.

